# Bacterial community structure analysis on *Listeria monocytogenes* inoculated spinach leaves is affected by PCR based methods to exclude chloroplast co-amplification

**DOI:** 10.1101/2024.02.01.578417

**Authors:** Paul Culliney, Achim Schmalenberger

## Abstract

Consumption of ready-to-eat leafy vegetables has increased in popularity due to their anticipated health benefits, but their consumption also poses a potential health risk in the form of foodborne pathogens. *Listeria monocytogenes* is a ubiquitous pathogen that has been regularly found on leafy vegetables including spinach. Growth determining factors go beyond plant species and cultivation practice and may include the phyllosphere bacteriome to affect the growth potential of *L. monocytogenes*. This study investigated the bacteriome of spinach leaves, stored under EURL challenge conditions for 9 days after inoculation with *L. monocytogenes* using two methods of excluding chloroplast co-amplification (COMPETE, BLOCK) at the PCR step as well as a post-PCR chloroplast sequence filter option (CONTROL). While all three approaches have demonstrated a change of bacterial communities over time, the pPNA based BLOCK approach resulted in greater diversity similarities to the CONTROL option. The COMPETE solution with a specifically designed primer to prevent chloroplast amplification had a strong underrepresentation of the Planctomycetota phylum and to a lesser extend underrepresentation of Chloroflexi and Verrucomicrobiota due to the inheritance of the selected primer region that allowed to deselect chloroplast co-amplification. However, the COMPETE approach had the lowest level of chloroplast co-amplification. Higher growth potential of *L. monocytogenes* from day 7 to 9 co-occurred with higher relative abundances of Pseudomonadaceae and lower numbers of Lactobacillales, suggesting that particular phylogenetic groups may support growth of *L. monocytogenes*. While chloroplast co-amplification with spinach in the present study was relatively modest and a purely filter based chloroplast removal was sufficient, other leafy vegetables may require one of the tested co-amplification prevention solutions. While the COMPETE solution in the present study was linked to some amplification bias, the approach may be useful when otherwise co-amplification is very high and the demonstrated BLOCK approach with pPNA is insufficient.

## Introduction

Plants such as leafy vegetables host diverse communities of microorganisms, collectively termed the microbiome (Dastogeer et al., 2020). These microorganisms inhabit either within (endosphere) or outer (epishere) surfaces of plant tissues. Moreover, the habitats of these microorganisms can be divided into above (phyllosphere) and below ground (rhizosphere). The phyllosphere is comprised of aerial parts of plants which for leafy vegetables is dominated by leaves. Global leaf surface area has been estimated at around 508,630,100 km^2^. Thus, when upper and lower leaf surface area are considered global leaf surface is 1,017,260,200 km^2^, which equates to roughly two times larger than global land surface area. Under the assumption that on average 10^6^ to 10^7^ microorganisms are present on each square cm of leaf surface, planetary epiphytic phyllosphere bacterial populations in excess of approximately 10^26^ could exist (Lindow and Brandl, 2003, Vorholt, 2012). Less is known about the characteristics and the ecological roles of the phyllosphere bacterial community members in comparison to the rhizosphere (Xu et al., 2022). Nonetheless, studies have investigated that the assembly of the phyllosphere microbiome is responsible for plant health and growth and are involved in protection for example by mediating plant-pathogen interactions (Daniell et al., 2016, Gong and Xin, 2021, Rastogi et al., 2013, Stone et al., 2018). Indeed, the utilisation of next generation sequencing (NGS) approaches such as Illumina sequencing have made it possible to conduct these types of research (Knief, 2014), A limitation of NGS of phyllosphere communities includes the co-amplification of mitochondrial and plastid (chloroplast) contamination as plant cells consist of DNA originating from its nucleus and two sub-cellular organelles i.e., the mitochondria and chloroplast (Gao et al., 2013, Hanshew et al., 2013). Chloroplasts are evolutionarily descended from bacteria; thus, their 16S gene sequences are almost identical to their bacterial ancestors. Therefore, primers which amplify bacterial 16S rRNA genes are also prone to co-amplify chloroplast SSU DNA (Sakai et al., 2004). Consequently, due to high abundance of plant DNA amplified, studies which do not prevent chloroplast DNA co-amplification could overestimate bacterial populations sizes in qPCR approaches (Rastogi et al., 2010) and affect bacterial community investigations. Indeed, using a PCR-DGGE approach to investigate the spinach phyllosphere resulted in a predominant band in the fingerprints that was identified as chloroplast (Lopez-Velasco et al., 2010). Likewise, next generation sequencing of the lettuce phyllosphere without chloroplast co-amplification supression resulted in around 90% of chloroplast amplicons, thus compromising reading depths of bacterial amplicon libraries (McManamon et al., 2019).

In efforts to block this co-amplification of chloroplast DNA, Peptide Nucleic Acid (PNAs) oligonuelcotides have been designed i.e., pPNA and mPNA for chloroplast and mitochondrial DNA, respectively. PNAs probes are artificially synthesized polymers of pyrimidine and purine nucelobases which are linked together by peptide bonds i.e., pseudo-peptide polymers as opposed to a deoxyribose phosphate backbone which is a distinct feature of DNA (Fitzpatrick et al., 2018). These oligonucleotides form thermally stable hybrid complexes by hybridising to complementary DNA sequences via a sequence dependent manner obeying typical Watson-Crick base pairing hydrogen bonding (Pellestor and Paulasova, 2004). Thus, due to the specifity of pPNA’s sequence (ggctcaaccctggacag) it has high binding affinity to chloroplast DNA of the selected target hypervariable region and blocks their amplification. pPNA clamps have been previously employed in research on plant based products such as strawberrry and tomato leaves, apple flower phytobiome and the root communities of 32 plant species (Fitzpatrick et al., 2018, Legein et al., 2022, Steven et al., 2018). Although, to our knowledge pPNA clamps have not been used to block the co-amplififcation of chloroplast DNA from the spinach phyllosphere. While pPNA clamps have proven to be highly effective at blocking chloroplast DNA of terrstertial and acquatic plants, this protocol has the potential to create a strong sequencing bias against Proteobacterial 16S rRNA (Jackrel et al., 2017). Utilising the common primer 784r which exploits a two base mismatch towards chloroplast DNA may reduce unwanted co-amplification. A recent study exploited this further by adding a competing chloroplast primer (784RInvT) with an inverted T at the 3-strand end of the oligo to prevent the polymerase from extending the oligonucleotide when bound to plant DNA (McManamon et al., 2019). McManamon and colleagues (2019) successfully utilised this method to prevent the extension of chloroplast DNA from iceberg lettuce and applied it to a nested PCR-DGGE fingerprint analysis.

The roles that phyllosphere bacteria play on plant leaves in general, and on leafy vegetables in particular, are still somewhat elusive. Bacterial communities may affect the growth of pathogens in their midst. Previous research has demonstrated that bacterial isolates from ready-to-eat (RTE) lettuce influence the survival of *Listeria* in co-cultures (Francis and O’Beirne, 2002). However, a paucity of studies have researched the *in-situ* influence of the food microbiome when it comes to growth of *L. monocytogenes*. More specifically, no studies have yet assessed the influence of the spinach phyllosphere community members on growth of *L. monocytogenes*. The determination of total bacterial count (TBC) of a food product is one of the supplementary tests conducted while undertaking growth potential challenge studies to determine whether that food can or cannot support *L. monocytogenes* growth (EURL Lm, 2021). However, the value of TBC determination is limited as it only enumerates the bacteria that can grow on the utilised standard medium which can be a minority of the total bacteria present on the food product. Therefore, cultivation independent analysis based on universal bacterial maker genes i.e., 16S rDNA is nowadays utilised to describe the bacterial community through 16S rDNA-based amplicon sequencing, using various next generation platforms. This technique provides detailed descriptions of bacterial phylogenetic groups down to the family or genus level contained in the food matrix, provided that chloroplast co-amplification is not compromising the analysis. This approach will allow to assess the presence or absence and relative abundance of bacteria important to the growth of *L. monocytogenes* on the spinach phyllosphere and the associated *L. monocytogenes* populations.

At the outset of this study, the hypothesis was that a chloroplast co-amplifcation suppression method is beneficial for accurate bacteriome analysis on spinach leaves and this could be used to better study the growth of *L. monocytogenes* in the bacterial community. Here, the pPNA based method (BLOCK) was tested alongside the solution that utilizes the competing primer for chloroplast DNA 784RInvT (COMPETE). The findings were compared to the bacteriome analysis without a blocking method only utilizing software based filtering tools post Illumina next generation sequencing. Therefore, the objectives of this study were to i) compare effectiveness of methods in reducing co-amplification of chloroplast DNA during 16S rRNA gene fragment amplification of prokaryotic DNA of a *L. monocytogenes* inoculated spinach phyllosphere, ii) report on their subsequent influence on bacterial community structures, iii) describe the progression of spinach’s phyllosphere community over time and iv) relate chloroplast DNA co-extraction and the associated phyllosphere community with growth of *L. monocytogenes*.

## Materials and methods

### 2.1 Extraction of DNA and DNA quantification

DNA extractions were conducted on 20 samples from *L. monocytogenes* growth potential experiments on open-field spinach (F1 Trumpet) that was stored for 0, 2, 5,7 and 9 days at 7°C for days 0-6 and at 12°C for days 7-9, where *L. monocytogenes* and total heterotrophic bacteria were enumerated on cultivation media (Culliney and Schmalenberger, 2022). Growth experiments were executed according to the EU guidance document’s guidelines for conducting growth potential studies (EURL Lm, 2019). Each of the 20 packages contained 25 g produce inoculated with 100 cfu g^-1^ of a three-strain mix of *L. monocytogenes* i.e., 959 (vegetable isolate), 1382 (EUR *Lm* reference strain), and 6179 (food processing plant isolate). Contents of each were transferred into separate stomacher bags and homogenized in 25 mL of PBS using a stomacher (Seward 400, AGB Scientific, Dublin, Ireland), for 120 s at a high speed (260 rpm) which were used for microbial analysis i.e., determination of *L. monocytogenes.* The associated average *L. monocytogenes* counts (log_10_ cfu g^-1^) across the five timepoints (+ or – the relative increase or decrease from the previous timepoint) were day 0 = 1.99, day 2 = 2.31 (+ 0.32), day 5 = 2.90 (+ 0.59), day 7 = 3.48 (+ 0.58), and day 9 = 4.58 (+ 1.10) (Culliney and Schmalenberger, 2022). The remaining obtained homogenate suspensions were transferred into 50 ml conical tubes and centrifuged at 4500 g (15 minutes at 4 °C). Supernatants were discarded and derived pellets were stored at −20 °C. For DNA extraction, pellets were resuspended with 400 μL of phosphate buffered saline (PBS) and 100 μL was utilized for DNA extraction with the PowerFood DNA Isolation kit (MO BIO Laboratories, Carlsbad, CA) according to manufacturer’s instructions. Quantification of the extracted DNA was conducted spectrophotometrically with the Take 3 plate in an Eon plate reader/incubator (BioTek, Winooski, VT).

### 2.2 16S rRNA gene amplification

In anticipation of significant co-amplification of plant DNA in the PCR (chloroplast 16S rRNA genes) as experienced previously with lettuce (McManamon et al., 2019), two methods were adopted to suppress amplification of plant DNA. Each of the three PCR protocols within this section were applied to the same 20 open-field spinach (F1 Trumpet) DNA samples as described above (2.1 Extraction of DNA and DNA quantification).

The first PCR protocol included primers 101 F (5′ACTGGCGGACGGGTGAGTAA’3) and 784R (5′TACCMGGGTATCTAATCCKG’3) targeting the V2-V4 regions to amplify preferentially 16S rRNA gene fragments from bacteria. Additionally, a third <u>oligonucleotide</u> was added to the reaction (784RinvT; 5′TACTGGGGTATCTAATCCCA’3T’5) which was optimised to bind to chloroplast DNA (McManamon et al., 2019). An inverted T at the 3-strand end of the oligo prevented the polymerase from extending the oligonucleotide when bound to plant DNA (COMPETE). Each 50 μl reaction contained 1 x buffer (2 mM MgCl_2_), 0.2 mM dNTP mix, 0.4 mmol of each primer (incl. 784RinvT), 0.5 U of DreamTaq polymerase (Fisher Scientific, Waltham, MA) and 1 μl template DNA (McManamon et al., 2019). The second PCR protocol (50 μl reaction) selected the V1-V9 region of the 16S rRNA genes using 0.4 mmol of each of the following primers 27F (5′AGAGTTTGATCMTGGCTCAG’3), 1492R (5′TACGGYTACCTTGTTACGACTT’3) and a pPNA probe_(5’GGCTCAACCCTGGACAG’3). PNA oligos were used as sequence-specific PCR blockers because PNA probes have strong binding affinity and specificity to its target DNA and not being recognized by DNA polymerase as a primer (Fitzpatrick et al., 2018). The pPNA probe used in this study blocked amplification of chloroplast 16S sequences from diverse plant species (BLOCK). The third PCR protocol (50 μl reaction) acted as a control amplifying the V1-V9 region using 0.4 mmol of primers 27F and 1492R, this time without any blocking of chloroplast (CONTROL).

For each of the 3 PCR protocols (COMPETE, BLOCK, CONTROL), amplification was conducted in a G-Storm cycler with the following PCR conditions: an initial denaturation step at 94 °C for 4 minutes, followed by 28 cycles of: denaturation at 94 °C for 45 seconds, annealing at 58°C for 45 seconds and extension at 72 °C for 60 seconds. Then lastly, one final cycle at 72 °C for 5 minutes. To confirm PCR products were amplified, PCR products were run on an 1.5 % agarose gel after each protocol.

### 2.3 Adding Illumina adaptors to amplicons for Next Generation Sequencing

PCR products obtained from the first round of amplification (COMPETE, BLOCK, CONTROL, see section 2.2) were purified using the GenElute™ PCR Clean-Up Kit (Sigma Aldrich, St. Louis, MO). The purified DNA was diluted 10-fold with DNA free sterile water and used as a DNA template for a nested PCR to attach Illumina adaptors for Illumina sequencing. The 50 µL PCR reaction included: 25 µL of 2X Kapa Hifi HS readymix, 1.5 mmol of primers 515F (5’TCGTCGGCAGCGTCAGATGTGTATAAGAGACAGGTGCCAGCMGCCGCGGTAA-3’) and 806R (5’GTCTCGTGGGCTCGGAGATGTGTATAAGAGACAGGGACTACHVGGGTWTCTAAT-3’) targeting the hypervariable region V3-V4 of the 16S rRNA gene and 1 µL of the DNA template. Conducted in a G-Storm cycler, PCR conditions were as follows: an initial denaturation step at 95 °C for 3 minutes, followed by 25 cycles of denaturation at 98 °C for 20 seconds, annealing at 65 °C for 15 seconds and extension at 72 °C for 15 seconds. To conclude, one final cycle was performed at 72 °C for one minute. PCR purification was conducted using GenElute™ PCR Clean-Up Kit (Sigma Aldrich) following manufacturer’s instructions. DNA analysis using the Take 3 plate and Eon plate reader (BioTek) revealed quantities of at least 26.4 ng µl^-1^ and a purity of at least 1.8 260/280nm.

### 2.4 Next Generation Sequencing

All 60 samples were sent to the University of Minnesota Genomics Center (UMGC) for indexing and Illumina MiSeq sequencing. Bioinformatics analysis was performed using QIIME2 2021.11 (Bolyen et al., 2019). Paired end sequences with quality were demultiplexed and imported with metadata via ManifestPhred33V2 file. This was followed by trimming and truncating (quality filtering at Q30) using the q2-dada2 plugin (Supplementary information: Table S1: Dada2 statistics). Assigning taxonomic information to the ASV sequences was conducted using a pre-trained Naive Bayes taxonomic classifier which was trained on the Silva version 138 99 % reference data set where sequences were trimmed to represent only the region between the 515F / 806R primers (V3V4 region). Sequences not assigned to a phylum level and mitochondrial sequences were removed using the filter-table method in the q2-taxa plugin. Following this, bioinformatics analysis was continued with and without chloroplast filtering using the previous filter-table method. Even sampling depths for use in diversity metrics were identified for chloroplast filtered and non-filtered groups. For chloroplast COMPETE, BLOCK and CONTROL group of 60 samples, an even sampling depth of 16,991 was applied. Rarefaction at that specified sampling depth retained 1,019,460 (33.27 %) features in 60 (100 %) samples. Whereas, when chloroplast sequences were not filtered from the same 60 samples, 19,837 was the sampling depth and 1,190,220 (37.46 %) features in 60 (100 %) samples were retained. These were confirmed as appropriate using alpha and beta rarefaction plots, as well as Spearman’s heat map. Alpha diversity metrics (observed ASVs, Shannon index, Pielou’s evenness and Faith’s Phylogenetic Diversity) and beta diversity metrics (unweighted UniFrac (Lozupone and Knight, 2005), weighted UniFrac (Lozupone et al., 2007), and Bray–Curtis dissimilarity) using q2-diversity were estimated with and without rarefaction and with and without chloroplast sequence filtering. (Supplementary materials; Tables S2 to S5). Alpha metrics were computed and visualised on boxplots and beta visualisations were computed and viewed on PCoA Emperor plots ). Analysis of Composition of Microbiomes (ANCOM) test in the q2-composition plugin was used to identify differentially abundant features i.e., identify individual taxa whose relative abundances are significantly different across the three different blocking protocols. Relative abundance was calculated in excel, after conversion of relative collapsed frequency biom tables (phylum and family levels) from QIIME2 to tsv files.

### 2.5 Statistical analysis

R-Studio software (version 4.1.1) was used for statistical analysis. In situations of normality (Shapiro-Wilk) and homoscedasticity (Levene’s) a one-way ANOVA was conducted to compare input, filtered, denoised, merged and non-chimeric reads between the three groups. The remainder of statistical analysis for alpha and beta diversity metrics was conducted in QIIME2. For alpha diversity (observed ASVs, Shannon index, Pielou’s evenness and Faith’s Phylogenetic Diversity (Faith, 1992)) comparisons between the three groups and pairwise comparisons were conducted through Kruskal-Wallis tests. Beta diversity was analysed through non-parametric permutation tests (999 permutations): PERMANOVA, ANOSIM and PERMDISP (Anderson and Walsh, 2013). Statistical significance was tested at P < 0.05. In situations of normality (Shapiro-Wilk) and homoscedasticity (Levene’s) a one-way ANOVA Tukey HSD post-hoc test applying Benjamini-Hochberg correction for multiple testing was conducted to compare relative abundances for all alpha diversity metrics and relative abundances across subgroups. In situations of non-normality, Kruskal-Wallis rank sum test with the function kruskal.test and Dunn test post-hoc analysis for multiple pairwise comparisons between groups was conducted, applying Benjamini-Hochberg correction for multiple testing. In situations of unequal variance, the function oneway.test was employed with var = F, and Games-Howell post-hoc analysis.

## Results

### 3.1 Dada2 statistics, mitochondria and chloroplast filtering using QIIME2

There were no significant differences identified between all three groups (ANOVA, p > 0.05) for input reads, filtered reads, denoised reads, merged reads, and non-chimeric reads (Supplementary Information: Table S1). On average, 50, 7 and 4 reads were removed from the COMPETE, BLOCK and CONTROL groups due to not being assigned a phylum classification level. The average number of mitochondrial reads discarded from complete 16S analysis from were 2,892, 1,341 and 844, respectively. From COMPETE, BLOCK and CONTROL, filtering of chloroplast sequences through QIIME2 plug-in removed on average 33, 146 and 5,475 reads per sample, respectively. Prior to exclusion using QIIME2, the number of chloroplast reads per sample in COMPETE ranged from 0 to 129 reads. For BLOCK and CONTROL the number reads present per sample ranged from 0 to 474 reads, and 642 to 13069, respectively.

### 3.2 Implications of chloroplast blocking method and time on alpha diversity

High levels of diversity were revealed by the alpha diversity metrics. Evidently, choice of PCR based chloroplast exclusion (COMPETE, BLOCK) or employing no blocking protocol (CONTROL) all led to differences amongst alpha diversities metrics (Supplementary information: Description S1 and Tables S2 to S5). Moreover, CONTROL had more similarities to both chloroplast exclusion groups, compared to COMPETE and BLOCK amongst themselves. While rarefaction did not greatly influence alpha diversity metrics, filtering out chloroplast reads in certain situations influenced level of significance and outcome of comparisons between groups. Therefore, this is a crucial step for the CONTROL where no exclusion of chloroplast amplification in the PCR step was applied.

Linear mixed effects models, irrespective of presence of chloroplast and whether rarefaction was applied or not, revealed that overall grouped comparisons (COMPETE, BLOCK and CONTROL) of observed features, Faith’s phylogenetic diversity, Shannon entropy and Pielou’s evenness were not significantly different across the five data points (Supplementary Information: Table S6). However, the appropriate statistical tests comparing each group separately across the different datapoints (day 0 to 9) revealed different outcomes (Supplementary Information: Tables S2 to S5). When chloroplast was both present and excluded, without rarefaction the following results were revealed: COMPETE group’s observed features and Faith’s phylogenetic diversity were overall significantly different although post-hoc tests did only reveal any significant differences between the datapoints day 2 to 9 for Faith’s phylogenetic diversity. For Shannon entropy and Pielou’s evenness, there was no overall significant difference. An overall significant difference was noted for BLOCK group’s observed features and Faith’ phylogenetic diversity, and post-hoc tests showed differences from day 0 to 9, 2 to 5, 2 to 7, 2 to 9, and 7 to 9. For Shannon entropy, the same outcome was identified without the post-hoc difference from day 7 to 9. While there was an overall significant difference for Pielou’s evenness for the BLOCK group, there was only a post-hoc difference from day 0 to 9. For the CONTROL group’s observed features, Faith’s phylogenetic diversity, and Pielou’s evenness no overall or post-hoc significant differences were revealed. Although, for control group’s Shannon entropy there was an overall significant difference with post-hoc identifying a significant difference from day 0 to 2. Thus, alpha diversity i.e., richness, biodiversity, abundance, and evenness of the taxa findings dependent on the chloroplast blocking method i.e., amongst groups but also within groups across time. While removing chloroplast did not influence the alpha diversity metrics across the different data points, the application of rarefaction did influence some of the post-hoc significance outcomes (Supplementary Information; Tables S2 to S5).

### 3.3 Implications of chloroplast blocking method and time on beta diversity

Principal coordinate analysis (PCoA, unweighted UniFrac distance matrix) revealed a clear separation between both BLOCK **(Figure 1; blue line)** and CONTROL **(red)** from the COMPETE **(orange)** group. This was confirmed with beta-diversity metrics (PERMANOVA p = 0.002; 0.002). There was no separation between BLOCK and CONTROL and no significant differences were identified (PERMANOVA p = 0.482). These observations remained the same irrespective of whether chloroplast reads were filtered out using QIIME2 (PERMANOVA p = 0.002, 0.002 and 0.193, respectively). Indeed, BLOCK and CONTROL group were most identical out of all groups with an unweighted UniFrac distance of 0.45. The remaining groups (COMPETE and CONTROL; BLOCK and COMPETE) were between 0.50 and 0.60 suggesting that those samples shared less than half of their branch length on the phylogenetic tree.

**Figure 1:**
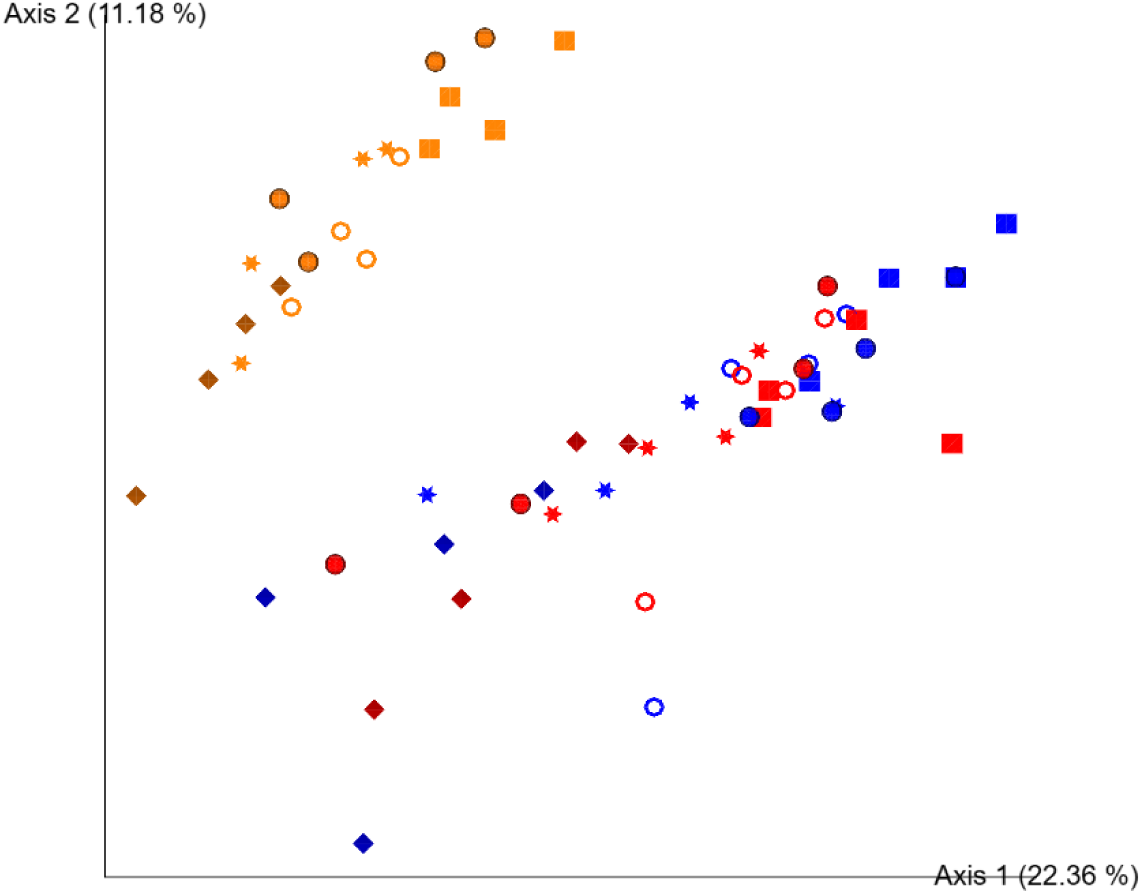
Two-dimensional Emperor (PCoA) plots showing beta diversity distances i.e., unweighted UniFrac, among the different samples across COMPETE (orange), BLOCK (blue) and CONTROL (red) groups with chloroplast reads filtered using Qiime2 plug-ins, with rarefaction applied. Shapes revealed separations across time where: day 0 = circle, day 2 = square, day 5 = star, day 7 = ring and day 9 = diamond.

However, when weighted according to proportional abundance, principal coordinate analysis **(Figure 2, PCoA, weighted UniFrac distance matrix)** revealed a clear separation between all three groups and significant differences were identified between all three (PERMANOVA; pairwise comparisons; p = 0.001) when chloroplast were not filtered out. Weighted UniFrac distances were between 0.10 to 0.15 for all group comparisons. Thus, by considering proportional abundance, it was revealed groups’ communities were even more similar and rare taxa were present within each of the groups which greatly inflated unweighted UniFrac distances. Filtering of chloroplast reads using QIIME2 had a substantial effect on the level of significance between weighted UniFrac distances of all three groups. In contrast, when chloroplast reads were removed post sequencing using QIIME2, no visible separation occurred between the BLOCK and CONTROL groups **(Figure 3 and 4; PCoA, weighted UniFrac distance matrix and Bray-Curtis)**. Although an overall significant difference was observed between all three groups, pairwise comparisons identified that BLOCK and CONTROL groups were not significantly different (PERMANOVA; weighted UniFrac and Bray-Curtis; p = 0.531 and 0.932). Moreover, separation was more evident and greater variation was explained by the PCoA plot for weighted UniFrac which considers phylogenetic relatedness compared to Bray-Curtis which does not. According to these data, BLOCK and CONTROL as well as BLOCK and COMPETE’s bacterial communities overlap more when compared to that of CONTROL and COMPETE, which is also reflected in the weighted UniFrac distances. Thus, filtering chloroplast sequences from CONTROL samples with no prior PCR based exclusion attempts was equally as effective as using the pPNA clamp in the present case of spinach phyllosphere analysis. In this study, rarefaction did not influence beta-diversity visualisations or metrics.

**Figure 2:**
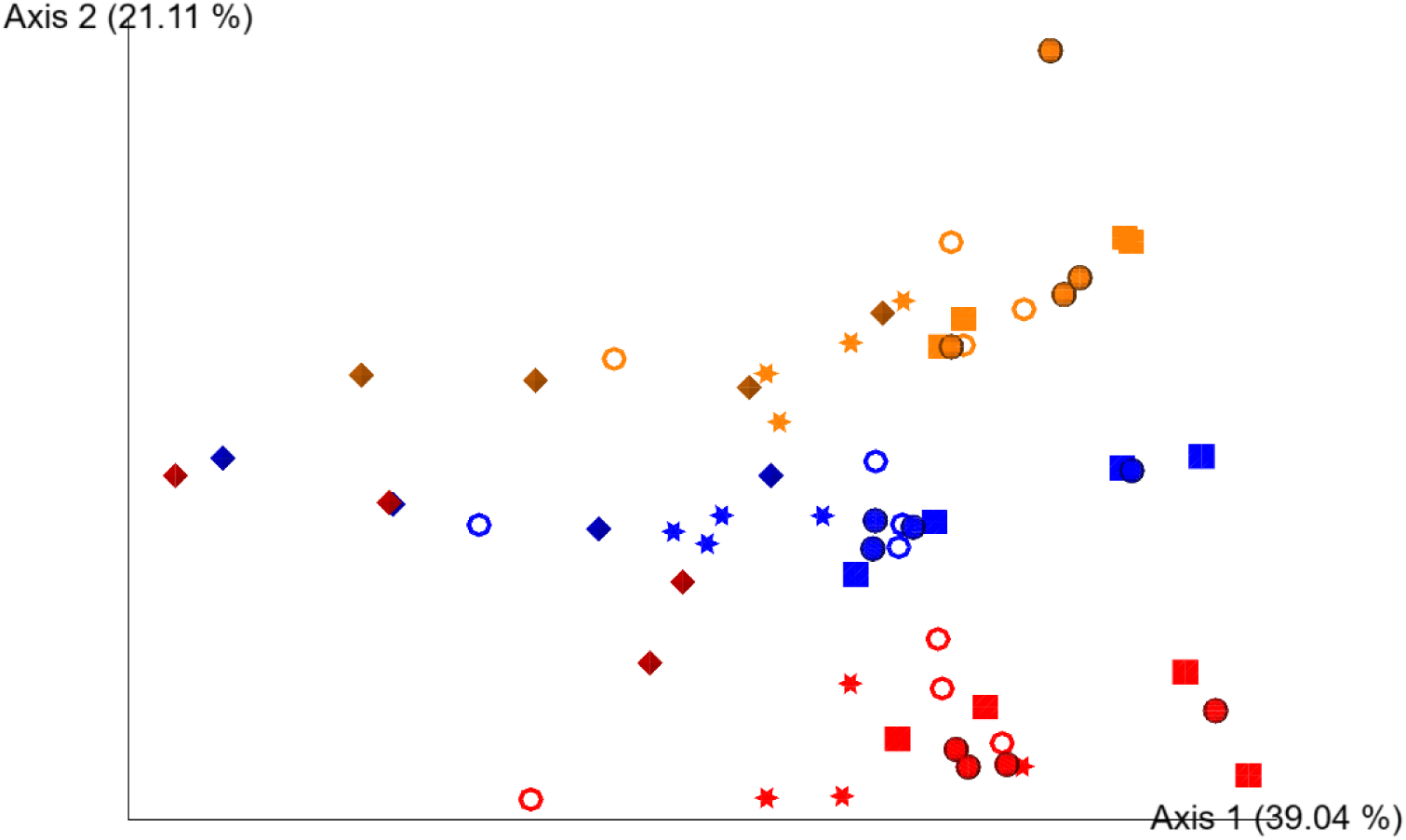
Two-dimensional Emperor (PCoA) plots showing beta diversity distances i.e., weighted UniFrac, among the different samples across COMPETE (orange), BLOCK (blue) and CONTROL (red) groups. Chloroplast reads were not filtered out using Qiime2 plug-ins, with rarefaction applied. Shapes revealed separations across time where: day 0 = circle, day 2 = square, day 5 = star, day 7 = ring and day 9 = diamond.

**Figure 3:**
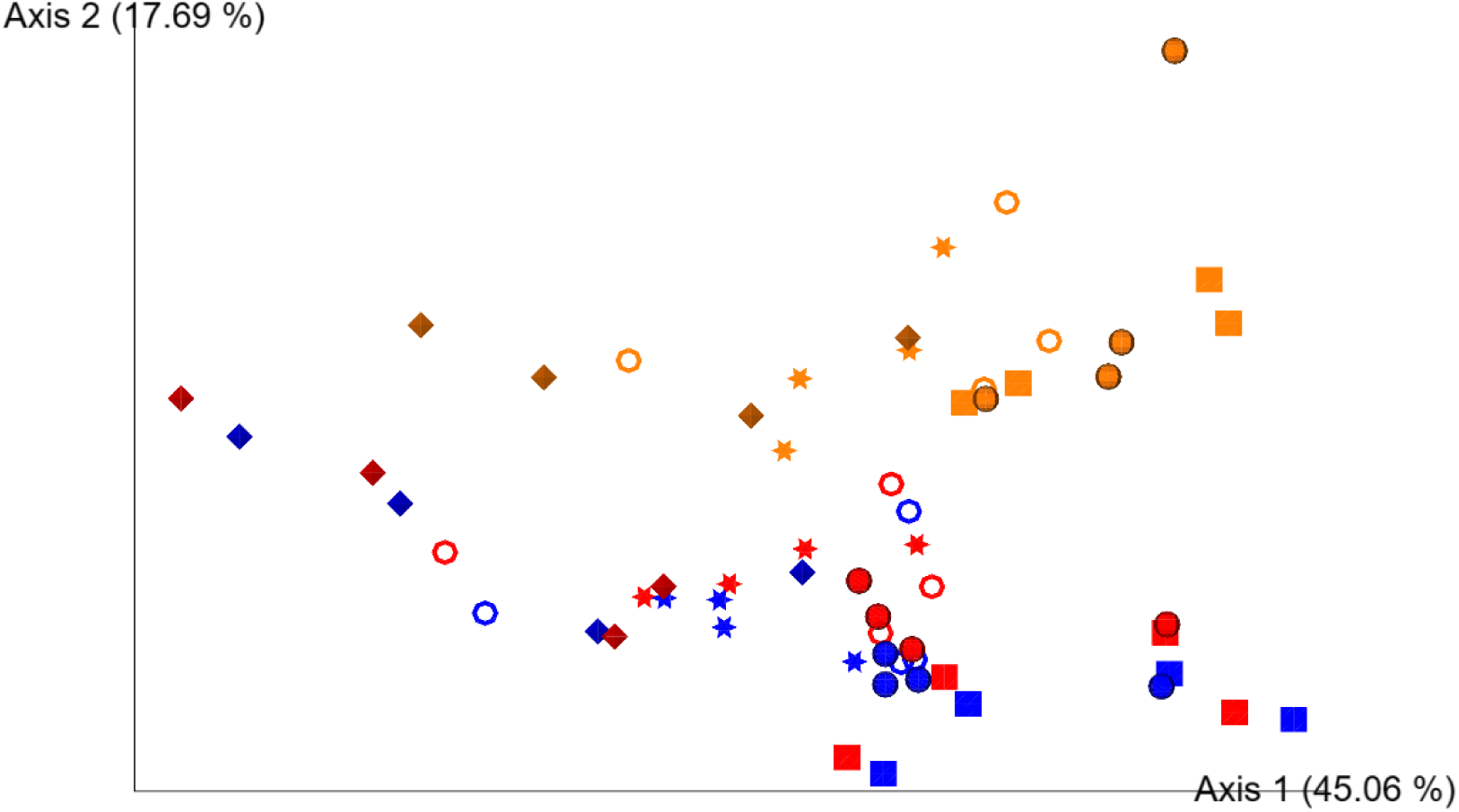
Two-dimensional Emperor (PCoA) plots showing beta diversity distances i.e., weighted UniFrac, among the different samples across COMPETE (orange), BLOCK (blue) and CONTROL (red) groups where chloroplast reads were filtered out using Qiime2 plug-ins, with rarefaction applied. Shapes revealed separations across time where: day 0 = circle, day 2 = square, day 5 = star, day 7 = ring and day 9 = diamond.

**Figure 4:**
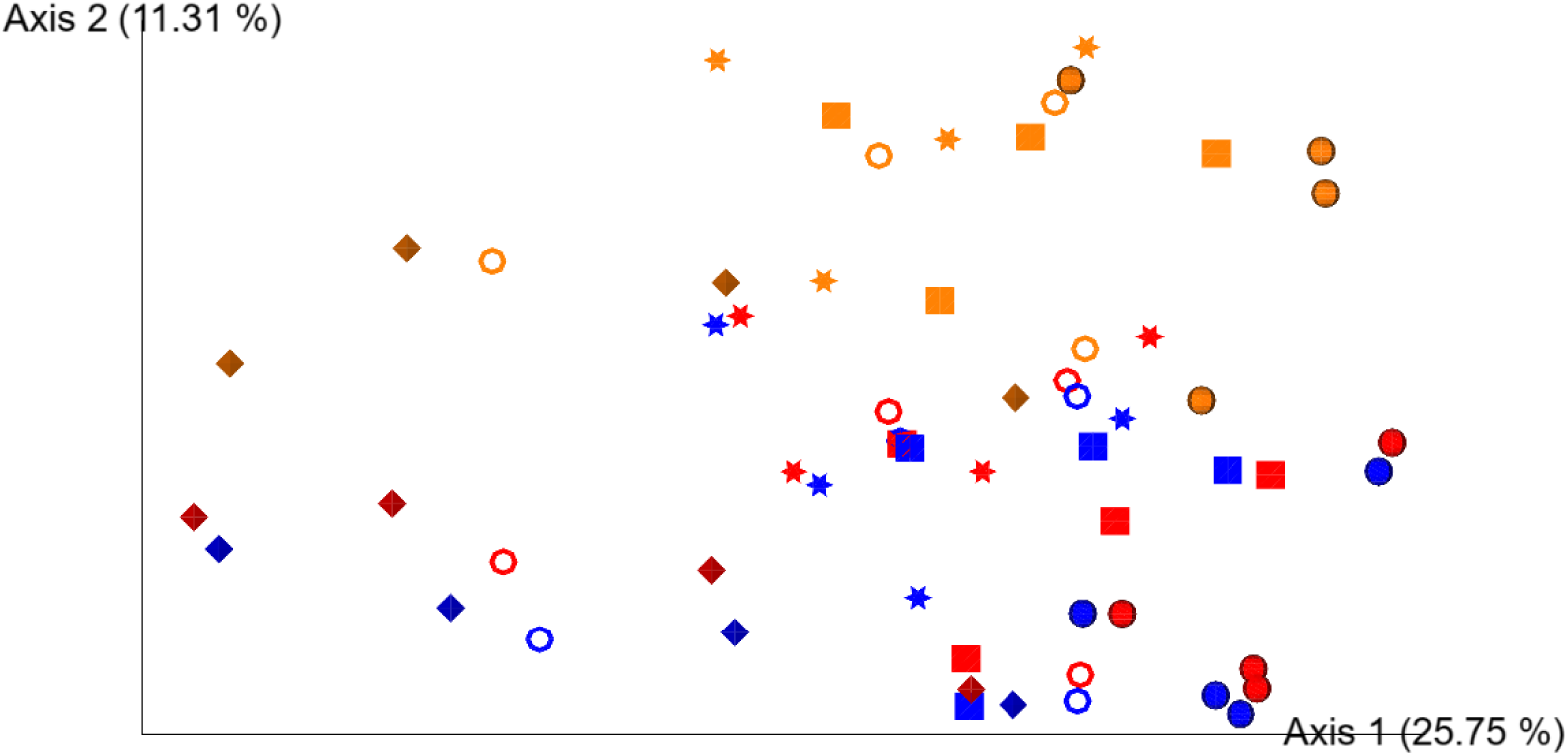
Two-dimensional Emperor (PCoA) plots showing beta diversity distances i.e., Bray Curtis, among the different samples across COMPETE (orange), BLOCK (blue) and CONTROL (red) groups where chloroplast reads were filtered out using Qiime2 plug-ins, with rarefaction applied. Shapes revealed separations across time where: day 0 = circle, day 2 = square, day 5 = star, day 7 = ring and day 9 = diamond.

A clear separation was observed of day 0, 2 and 7 (circle, square and ring) from day 5 (star) with a further separation from day 9 (diamond) **(Figure 3; PCoA, weighted UniFrac distance matrix)**. These separations were accompanied by significant differences across time for each group. For COMPETE, significant separations were identified for day 0 to 5 (p = 0.038), day 0 to 9 (p = 0.031), day 2 to 5 (p = 0.031) and day 2 to 9 (p = 0.039). An additional significant separation was observed for BLOCK, i.e., day 0 to 5 (p = 0.029), day 0 to 9 (p = 0.031), day 2 to 5 (p = 0.032), day 2 to 7 (p = 0.018) and day 2 to 9 (p = 0.033). Fewer significant separations were identified for CONTROL, i.e., day 0 to 5 (p = 0.025), day 0 to 9 (p = 0.023) and day 2 to 9 (p = 0.030). In either case, all groups collectively displayed changes in bacterial communities over time with significant separations from day 0 to 5, day 0 to 9 and day 2 to 9. Moreover, comparisons between groups across time also had significant separations based on weighted UniFrac distances. COMPETE and BLOCK were significantly separated at day 0 (p = 0.025), day 2 (p = 0.034) and day 5 (p = 0.026). COMPETE and CONTROL were significantly separated at day 0 (p = 0.035) and day 2 (p = 0.032). As observed in **Figure 3**, there were no significant separations across time between BLOCK and CONTROL (p = 0.507 to 0.834).

### 3.4 Effectiveness of chloroplast blocking protocols

Chloroplast relative abundance on average was 10.93 % for CONTROL. The pPNA clamp (BLOCK) was highly effective at blocking the co-amplification of chloroplast DNA with an average abundance of 0.26 % and therefore achieved a 40-fold reduction in the co-amplification of chloroplast DNA. The chloroplast content of all three groups were all significantly different **(p < 0.05; Table 1)**. Therefore, the most effective of the PCR protocols at blocking chloroplast co-amplification was the RInvT primer method (COMPETE) with a relative abundance of 0.06 % representing a 180-fold reduction in chloroplast co-amplification. COMPETE was four times more efficient in reducing chloroplast co-amplification compared to BLOCK. Moreover, as almost all chloroplast co-amplification was prevented, the chloroplast contents remained stable for COMPETE over time whereas, a significant reduction over time was identified for BLOCK. A similar significant reduction in chloroplast content at day 9 was identified for CONTROL, where chloroplast contents varied between 10 and 16 % over days 0 to 7 and then dropped to below 3 % at day 9.

**Table 1:** Chloroplast content of each group across time prior to removal using Qiime2. ± standard error. Letters a – d indicate signifficance within groups. Letters A – B indicate significance between groups.

| Removal<br>method | Day 0 | Day 2 | Day 5 | Day 7 | Day 9 |  |
| --- | --- | --- | --- | --- | --- | --- |
| **COMPETE** | 0.09% ± 0.02<sup>a</sup> | 0.09% ± 0.01<sup>a</sup> | 0.08% ± 0.04<sup>a</sup> | 0.03% ± 0.01<sup>a</sup> | 0.00% ± 0.00<sup>a</sup> | **A** |
| **BLOCK** | 0.24% ± 0.02<sup>ab</sup> | 0.31% ± 0.14<sup>ab</sup> | 0.48% ± 0.17<sup>a</sup> | 0.25% ± 0.05<sup>ab</sup> | 0.03% ± 0.02<sup>b</sup> | **B** |
| **CONTROL** | 14.04% ± 0.38<sup>a</sup> | 10.38% ± 1.28<sup>a</sup> | 16.29% ± 3.01<sup>a</sup> | 11.33% ± 0.92<sup>a</sup> | 2.64% ± 0.95<sup>b</sup> | **C** |

### 3.5 Relative abundance of phyla and families amongst groups and across time

The top four most abundant phyla for all groups were Proteobacteria, Bacteroidota, Actinobacteriota, and Firmicutes **(Figure 5)**. These four phyla represented 96.32, 94.50 and 95.25 % of total phyla of the COMPETE, BLOCK and CONTROL groups, respectively. Across time for COMPETE these remained the four most abundant phyla and their total abundance ranged from 94.28 to 98.24 % of total phyla. Similarly, across time for BLOCK and CONTROL, these remained the four most abundant phyla and their total abundance ranged from 89.30 to 98.78 % and 90.67 to 98.25 % of total phyla, respectively.

**Figure 5:**
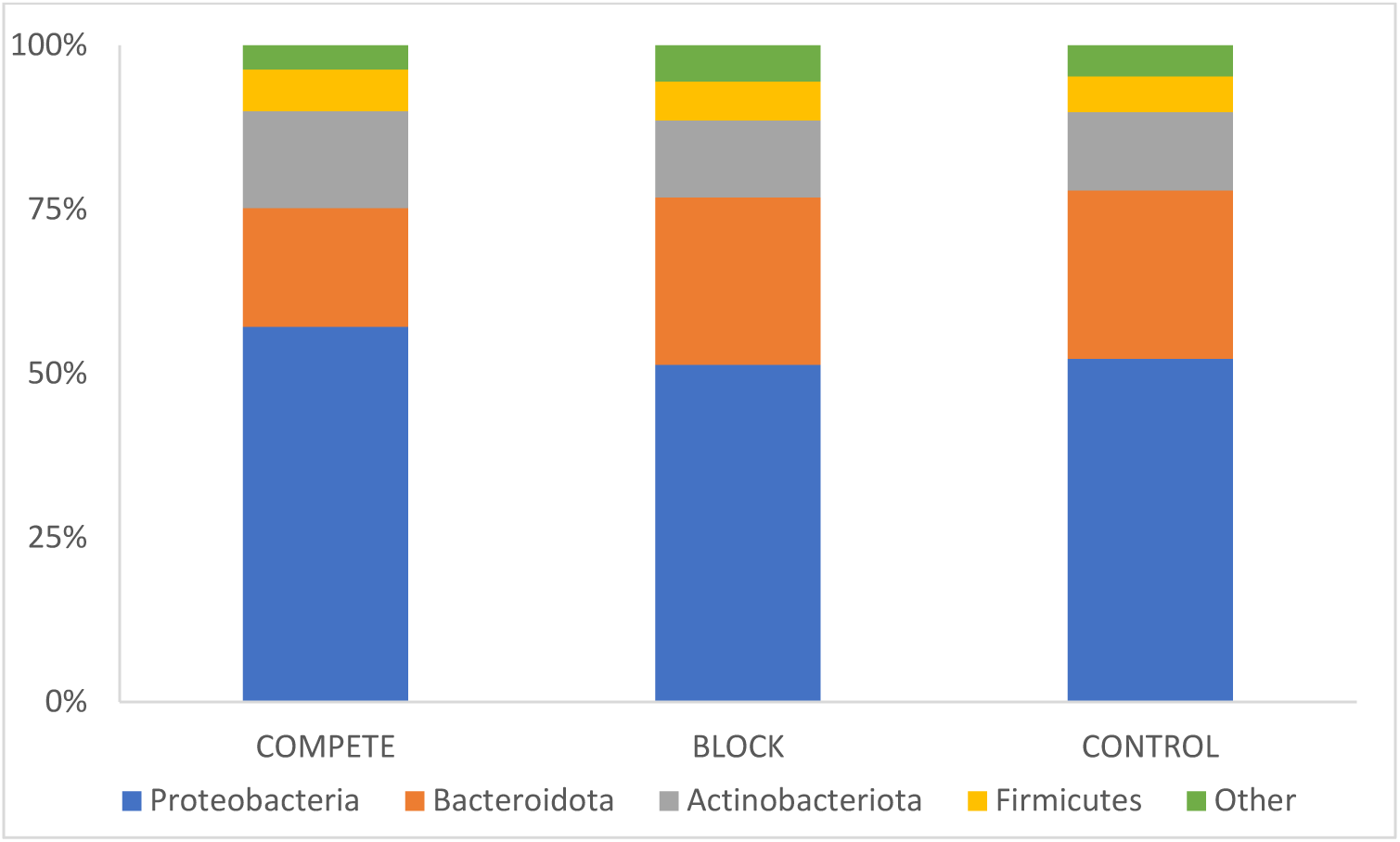
Mean relative abundances (%) of four most abundant phyla of the 16S gene of the COMPETE, BLOCK and CONTROL groups, with chloroplast removed using Qiime2 software and rarefaction applied. All remaining lower abundant phyla are combined in “Other”.

Of the 20 significantly different families identified in **Table 2** (any group with a family relative abundance > 0.20 %) between the three groups, 16 families were associated with significant pairwise differences between COMPETE and both BLOCK and CONTROL, but not between BLOCK and CONTROL themselves. Out of the top 20 most abundant families for each group, 17 families were shared by COMPETE, BLOCK and CONTROL **(Table 3 to 5)**. For the COMPETE group overall significant differences (p < 0.05) across time were revealed for Pseudomonadaceae, Nocardiaceae, Sphingobacteriaceae, Beijerinckiaceae, Microbacteriaceae and Flavobacteriaceae. For the BLOCK group, this occurred for Pseudomonadaceae, Hymenobacteraceae, Sphingobacteriaceae, Beijerinckiaceae, Rhizobioceae, Flavobacteriaceae and Pirellulaceae. Finally, for the CONTROL group, significant differences across time were identified for Pseudomonadaceae, Sphingomonadaceae, Nocardiaceae, Hymenobacteraceae, Sphingobacteriaceae, Beijerinckiaceae and Flavobacteriaceae. Therefore, the bacterial community structures (based on 16S rRNA gene sequence) at family level were affected by the chloroplast blocking method selected and were influenced by the duration of the incubation. Although, from day 0 to 9, significantly increasing relative abundances of families Pseudomonadaceae, Sphingobacteriaceae, and Flavobacteriaceae and significantly decreasing relative abundances of Beijerinckiaceae were commonly associated with all three groups.

**Table 2:** Average relative abundance ± the standard error of families present in the phyllosphere of the COMPETE, BLOCK and CONTROL groups with chloroplast reads filtered using Qiime2 software, with rarefaction applied. Families listed are present with greater than 0.2 % relative abundance in at least one of the three group. All remaining lower abundant families are combined in “Other”. * indicates an overall significant difference for that family. a to b indicate significance differences between the groups.

| Families | COMPETE | BLOCK | CONTROL |
| --- | --- | --- | --- |
| Rubinisphaeraceae* | 0.00 ± 0.00<sup>a</sup> | 0.21 ± 0.05<sup>b</sup> | 0.16 ± 0.03<sup>b</sup> |
| Gemmataceae* | 0.00 ± 0.00<sup>a</sup> | 0.43 ± 0.10<sup>b</sup> | 0.30 ± 0.07<sup>b</sup> |
| Chthoniobacteraceae* | 0.07 ± 0.01<sup>a</sup> | 0.39 ± 0.06<sup>b</sup> | 0.56 ± 0.12<sup>b</sup> |
| Spirosomaceae* | 0.36 ± 0.04<sup>a</sup> | 1.01 ± 0.10<sup>b</sup> | 1.01 ± 0.10<sup>b</sup> |
| Pseudomonadaceae* | 12.89 ± 0.78<sup>a</sup> | 8.25 ± 0.84<sup>b</sup> | 9.17 ± 0.75<sup>b</sup> |
| Myxococcaceae* | 0.37 ± 0.04<sup>a</sup> | 0.11 ± 0.02<sup>b</sup> | 0.32 ± 0.04<sup>a</sup> |
| Hymenobacteraceae | 6.56 ± 0.74<sup>a</sup> | 6.61 ± 0.71<sup>a</sup> | 6.58 ± 0.63<sup>a</sup> |
| Sphingomonadaceae* | 8.70 ± 0.59<sup>a</sup> | 11.18 ± 0.57<sup>b</sup> | 11.92 ± 0.77<sup>b</sup> |
| Isosphaeraceae* | 0.00 ± 0.00<sup>a</sup> | 0.22 ± 0.05<sup>b</sup> | 0.11 ± 0.02<sup>b</sup> |
| Nocardiodiaceae* | 0.61 ± 0.05<sup>a</sup> | 0.22 ± 0.03<sup>b</sup> | 0.22 ± 0.03<sup>b</sup> |
| Pirellulaceae* | 0.00 ± 0.00<sup>a</sup> | 0.98 ± 0.19<sup>b</sup> | 0.64 ± 0.13<sup>b</sup> |
| Enterobacterales; unknown* | 0.53 ± 0.07<sup>a</sup> | 0.30 ± 0.05<sup>b</sup> | 0.43 ± 0.06<sup>ab</sup> |
| Sphingobacteriaceae | 6.28 ± 0.81<sup>a</sup> | 6.12 ± 0.82<sup>a</sup> | 6.69 ± 0.94<sup>a</sup> |
| Weeksellaceae* | 2.49 ± 0.43<sup>a</sup> | 8.72 ± 0.63<sup>b</sup> | 8.33 ± 0.52<sup>b</sup> |
| Uncultured* | 0.02 ± 0.01<sup>a</sup> | 0.28 ± 0.05<sup>b</sup> | 0.21 ± 0.05<sup>b</sup> |
| Xanthomonadaceae* | 3.04 ± 0.28<sup>a</sup> | 4.24 ± 0.35<sup>b</sup> | 3.23 ± 0.31<sup>a</sup> |
| Flavobacteriaceae | 2.37 ± 0.73<sup>a</sup> | 2.85 ± 0.81<sup>a</sup> | 2.84 ± 0.89<sup>a</sup> |
| Rhizobiaceae | 5.40 ± 0.38<sup>a</sup> | 5.04 ± 0.42<sup>a</sup> | 4.90 ± 0.33<sup>a</sup> |
| Caulobacteraceae | 0.96 ± 0.12<sup>a</sup> | 0.98 ± 0.16<sup>a</sup> | 0.78 ± 0.10<sup>a</sup> |
| Leuconostocaceae | 0.21 ± 0.11<sup>a</sup> | 0.17 ± 0.08<sup>a</sup> | 0.16 ± 0.08<sup>a</sup> |
| Microbacteriaceae | 5.00 ± 0.45<sup>a</sup> | 5.71 ± 0.34<sup>a</sup> | 5.49 ± 0.37<sup>a</sup> |
| Beijerinckiaceae* | 5.46 ± 0.53<sup>a</sup> | 4.34 ± 0.47<sup>a</sup> | 3.84 ± 0.38<sup>a</sup> |
| Nocardiaceae* | 7.35 ± 0.55<sup>a</sup> | 4.44 ± 0.47<sup>b</sup> | 5.10 ± 0.48<sup>b</sup> |
| Xanthobacteraceae | 0.15 ± 0.02<sup>a</sup> | 0.20 ± 0.04<sup>a</sup> | 0.20 ± 0.05<sup>a</sup> |
| Micrococcaceae | 0.77 ± 0.11<sup>a</sup> | 0.75 ± 0.09<sup>a</sup> | 0.61 ± 0.09<sup>a</sup> |
| Paenibacillaceae | 4.19 ± 0.24<sup>a</sup> | 3.84 ± 0.27<sup>a</sup> | 3.53 ± 0.25<sup>a</sup> |
| Bacillaceae | 0.28 ± 0.08<sup>a</sup> | 0.29 ± 0.08<sup>a</sup> | 0.31 ± 0.10<sup>a</sup> |
| Yersiniaceae | 1.11 ± 0.49<sup>a</sup> | 0.81 ± 0.40<sup>a</sup> | 0.97 ± 0.46<sup>a</sup> |
| Xiphinematobacteraceae* | 0.08 ± 0.01<sup>a</sup> | 0.24 ± 0.04<sup>b</sup> | 0.39 ± 0.08<sup>b</sup> |
| Moraxellaceae | 0.82 ± 0.18<sup>a</sup> | 0.52 ± 0.14<sup>a</sup> | 0.67 ± 0.15<sup>a</sup> |
| Oxalobacteraceae | 4.38 ± 0.22<sup>a</sup> | 4.26 ± 0.28<sup>a</sup> | 3.89 ± 0.20<sup>a</sup> |
| Comamonadaceae | 2.68 ± 0.21<sup>a</sup> | 2.80 ± 0.21<sup>a</sup> | 2.48 ± 0.18<sup>a</sup> |
| Deinococcaceae* | 1.20 ± 0.20<sup>a</sup> | 0.65 ± 0.09<sup>b</sup> | 0.42 ± 0.06<sup>b</sup> |
| Exiguobacteraceae | 1.30 ± 0.28<sup>a</sup> | 1.20 ± 0.29<sup>a</sup> | 1.10 ± 0.26<sup>a</sup> |
| Rhodanobacteraceae | 0.64 ± 0.17<sup>a</sup> | 0.77 ± 0.21<sup>a</sup> | 0.52 ± 0.12<sup>a</sup> |
| Saccharimonadaceae* | 1.25 ± 0.20<sup>a</sup> | 0.17 ± 0.04<sup>b</sup> | 0.28 ± 0.04<sup>b</sup> |
| Enterobacteriaceae | 2.53 ± 0.45<sup>a</sup> | 1.84 ± 0.32<sup>a</sup> | 2.24 ± 0.41<sup>a</sup> |
| Kineosporiaceae* | 0.21 ± 0.02<sup>a</sup> | 0.10 ± 0.01<sup>b</sup> | 0.09 ± 0.01<sup>b</sup> |
| Erwiniaceae | 7.00 ± 0.95<sup>a</sup> | 5.18 ± 0.84<sup>a</sup> | 6.41 ± 0.87<sup>a</sup> |
| Other | 2.74 | 3.58 | 2.9 |

**Table 3:** Average relative abundance (% ± standard error) of the 20 most abundant families (16S rDNA) of the COMPETE group across days 0, 2, 5. 7, and 9 (chloroplasts filtered with Qiime2, rarefied). * indicates an overall significant result was identified for that particular family. Letters a – e indicate significant differences over time. 17 common families between COMPETE, BLOCK and CONTROL groups are displayed first. All remaining lower abundant families are combined in “Other”.

| COMPETE | Day 0 | Day 2 | Day 5 | Day 7 | Day 9 |
| --- | --- | --- | --- | --- | --- |
| Pseudomonadaceae* | 12.63 ± 0.98<sup>ab</sup> | 9.24 ± 0.35<sup>a</sup> | 15.43 ± 0.94<sup>b</sup> | 12.40 ± 1.88<sup>ab</sup> | 14.74 ± 2.47<sup>ab</sup> |
| Sphingomonadaceae | 9.27 ± 2.15<sup>a</sup> | 9.31 ± 0.56<sup>a</sup> | 8.56 ± 1.14<sup>a</sup> | 8.96 ± 1.47<sup>a</sup> | 7.38 ± 1.39<sup>a</sup> |
| Nocardiaceae* | 6.04 ± 0.75<sup>a</sup> | 10.41 ± 1.32<sup>b</sup> | 7.26 ± 1.27<sup>ab</sup> | 7.22 ± 0.38<sup>ab</sup> | 5.80 ± 1.00<sup>a</sup> |
| Erwiniaceae | 11.10 ± 0.53<sup>a</sup> | 4.91 ± 1.64<sup>a</sup> | 7.17 ± 2.33<sup>a</sup> | 6.78 ± 3.02<sup>a</sup> | 5.04 ± 1.69<sup>a</sup> |
| Hymenobacteraceae | 9.50 ± 2.68<sup>a</sup> | 7.54 ± 0.61<sup>a</sup> | 6.95 ± 0.48<sup>a</sup> | 5.43 ± 0.97<sup>a</sup> | 3.39 ± 1.19<sup>a</sup> |
| Sphingobacteriaceae* | 2.33 ± 0.67<sup>a</sup> | 4.34 ± 1.23<sup>ab</sup> | 6.74 ± 1.09<sup>ab</sup> | 6.99 ± 1.50<sup>ab</sup> | 10.98 ± 1.21<sup>b</sup> |
| Beijerinckiaceae* | 7.53 ± 0.48<sup>a</sup> | 7.33 ± 1.05<sup>a</sup> | 5.22 ± 0.83<sup>ab</sup> | 4.63 ± 0.95<sup>ab</sup> | 2.57 ± 0.44<sup>b</sup> |
| Rhizobiaceae | 4.34 ± 1.08<sup>a</sup> | 6.44 ± 0.79<sup>a</sup> | 4.68 ± 0.64<sup>a</sup> | 4.63 ± 0.63<sup>a</sup> | 6.90 ± 0.42<sup>a</sup> |
| Microbacteriaceae* | 7.79 ± 0.33<sup>a</sup> | 4.59 ± 0.94<sup>b</sup> | 3.76 ± 0.17<sup>b</sup> | 4.47 ± 1.22<sup>b</sup> | 4.38 ± 0.66<sup>b</sup> |
| Oxalobacteraceae | 3.97 ± 0.40<sup>a</sup> | 4.45 ± 0.36<sup>a</sup> | 4.38 ± 0.53<sup>a</sup> | 4.04 ± 0.55<sup>a</sup> | 5.05 ± 0.68<sup>a</sup> |
| Paenibacillaceae | 4.40 ± 0.25<sup>a</sup> | 3.28 ± 0.19<sup>a</sup> | 3.79 ± 0.17<sup>a</sup> | 5.04 ± 0.71<sup>a</sup> | 4.42 ± 0.81<sup>a</sup> |
| Xanthomonadaceae | 2.14 ± 0.30<sup>a</sup> | 3.31 ± 0.96<sup>a</sup> | 2.76 ± 0.56<sup>a</sup> | 3.42 ± 0.55<sup>a</sup> | 3.57 ± 0.63<sup>a</sup> |
| Comamonadaceae | 3.11 ± 0.13<sup>a</sup> | 3.33 ± 0.07<sup>a</sup> | 2.07 ± 0.29<sup>a</sup> | 2.55 ± 0.89<sup>a</sup> | 2.31 ± 0.41<sup>a</sup> |
| Enterobacteriaceae | 1.32 ± 0.72<sup>a</sup> | 1.97 ± 0.63<sup>a</sup> | 2.54 ± 0.59<sup>a</sup> | 2.29 ± 0.85<sup>a</sup> | 4.53 ± 1.62<sup>a</sup> |
| Weeksellaceae | 1.09 ± 0.24<sup>a</sup> | 1.65 ± 0.78<sup>a</sup> | 4.06 ± 1.18<sup>a</sup> | 2.31 ± 0.80<sup>a</sup> | 3.35 ± 1.12<sup>a</sup> |
| Flavobacteriaceae* | 0.25 ± 0.07<sup>a</sup> | 0.44 ± 0.09<sup>ab</sup> | 2.76 ± 0.76<sup>ab</sup> | 2.13 ± 1.74<sup>ab</sup> | 6.29 ± 2.33<sup>b</sup> |
| Exiguobacteraceae | 1.52 ± 0.24<sup>a</sup> | 1.35 ± 0.21<sup>a</sup> | 0.89 ± 0.30<sup>a</sup> | 2.37 ± 1.20<sup>a</sup> | 0.37 ± 0.11<sup>a</sup> |
| Saccharimonadaceae | 0.16 ± 0.05<sup>a</sup> | 0.13 ± 0.07<sup>a</sup> | 0.04 ± 0.01<sup>a</sup> | 0.08 ± 0.01<sup>a</sup> | 0.01 ± 0.01<sup>a</sup> |
| Deinococcaceae | 2.05 ± 0.79<sup>a</sup> | 1.28 ± 0.36<sup>a</sup> | 0.93 ± 0.12<sup>a</sup> | 1.08 ± 0.42<sup>a</sup> | 0.66 ± 0.12<sup>a</sup> |
| Yersiniaceae | 0.26 ± 0.03<sup>a</sup> | 0.52 ± 0.07<sup>a</sup> | 1.01 ± 0.18<sup>a</sup> | 2.89 ± 2.45<sup>a</sup> | 0.87 ± 0.31<sup>a</sup> |
| Other | 9.20 | 14.18 | 9.00 | 10.29 | 7.39 |

**Table 4:** Average relative abundance (% ± standard error) of the 20 most abundant families (16S rDNA) of the BLOCK group across days 0, 2, 5. 7, and 9 (chloroplasts filtered with Qiime2, rarefied). * indicates an overall significant result was identified for that particular family. Letters a – e indicate significant differences over time. 17 common families between COMPETE, BLOCK and CONTROL groups are displayed first. All remaining lower abundant families are combined in “Other”.

| BLOCK | Day 0 | Day 2 | Day 5 | Day 7 | Day 9 |
| --- | --- | --- | --- | --- | --- |
| Pseudomonadaceae* | 6.80 ± 0.22<sup>ab</sup> | 4.37 ± 0.49<sup>a</sup> | 11.56 ± 1.82<sup>b</sup> | 8.36 ± 1.52<sup>ab</sup> | 10.16 ± 2.46<sup>ab</sup> |
| Sphingomonadaceae | 13.40 ± 0.71<sup>a</sup> | 11.05 ± 0.92<sup>a</sup> | 12.38 ± 1.41<sup>a</sup> | 10.28 ± 1.38<sup>a</sup> | 8.78 ± 0.84<sup>a</sup> |
| Nocardiaceae | 3.01 ± 0.29<sup>a</sup> | 7.00 ± 1.41<sup>a</sup> | 4.73 ± 1.03<sup>a</sup> | 3.96 ± 0.22<sup>a</sup> | 3.50 ± 0.80<sup>a</sup> |
| Erwiniaceae | 8.43 ± 0.58<sup>a</sup> | 3.32 ± 1.25<sup>a</sup> | 3.88 ± 1.28<sup>a</sup> | 6.44 ± 3.30<sup>a</sup> | 3.83 ± 1.15<sup>a</sup> |
| Hymenobacteraceae* | 10.64 ± 0.89<sup>a</sup> | 7.53 ± 0.72<sup>ab</sup> | 5.91 ± 1.25<sup>bc</sup> | 6.06 ± 1.27<sup>bc</sup> | 2.93 ± 1.00<sup>c</sup> |
| Sphingobacteriaceae* | 3.56 ± 0.68<sup>a</sup> | 3.50 ± 0.94<sup>a</sup> | 6.37 ± 1.26<sup>a</sup> | 6.20 ± 1.10<sup>a</sup> | 10.95 ± 2.25<sup>a</sup> |
| Beijerinckiaceae* | 5.20 ± 0.60<sup>ab</sup> | 6.55 ± 1.09<sup>a</sup> | 4.44 ± 1.07<sup>ab</sup> | 3.35 ± 0.64<sup>ab</sup> | 2.17 ± 0.49<sup>b</sup> |
| Rhizobiaceae* | 4.03 ± 0.91<sup>a</sup> | 5.41 ± 0.84<sup>a</sup> | 3.99 ± 0.73<sup>a</sup> | 4.48 ± 0.85<sup>a</sup> | 7.28 ± 0.53<sup>a</sup> |
| Microbacteriaceae | 6.72 ± 0.46<sup>a</sup> | 5.27 ± 0.43<sup>a</sup> | 5.41 ± 0.57<sup>a</sup> | 4.85 ± 0.81<sup>a</sup> | 6.29 ± 1.26<sup>a</sup> |
| Oxalobacteraceae | 4.77 ± 0.50<sup>a</sup> | 4.56 ± 0.30<sup>a</sup> | 4.51 ± 1.06<sup>a</sup> | 3.61 ± 0.60<sup>a</sup> | 3.83 ± 0.56<sup>a</sup> |
| Paenibacillaceae | 4.05 ± 0.35<sup>a</sup> | 2.92 ± 0.35<sup>a</sup> | 3.63 ± 0.41<sup>a</sup> | 4.80 ± 0.85<sup>a</sup> | 3.82 ± 0.72<sup>a</sup> |
| Xanthomonadaceae | 3.89 ± 0.77<sup>a</sup> | 4.23 ± 1.31<sup>a</sup> | 4.21 ± 0.83<sup>a</sup> | 4.48 ± 0.33<sup>a</sup> | 4.39 ± 0.87<sup>a</sup> |
| Comamonadaceae | 3.18 ± 0.14<sup>a</sup> | 3.65 ± 0.12<sup>a</sup> | 2.13 ± 0.56<sup>a</sup> | 2.74 ± 0.74<sup>a</sup> | 2.32 ± 0.21<sup>a</sup> |
| Enterobacteriaceae | 1.02 ± 0.43<sup>a</sup> | 1.42 ± 0.54<sup>a</sup> | 1.80 ± 0.67<sup>a</sup> | 1.92 ± 0.71<sup>a</sup> | 3.04 ± 1.00<sup>a</sup> |
| Weeksellaceae | 7.01 ± 1.30<sup>a</sup> | 7.00 ± 1.42<sup>a</sup> | 9.79 ± 1.33<sup>a</sup> | 8.35 ± 0.83<sup>a</sup> | 11.43 ± 1.28<sup>a</sup> |
| Flavobacteriaceae* | 0.37 ± 0.05<sup>a</sup> | 0.56 ± 0.16<sup>ab</sup> | 3.65 ± 0.76<sup>ab</sup> | 2.82 ± 2.18<sup>ab</sup> | 6.84 ± 2.47<sup>b</sup> |
| Exiguobacteraceae | 0.99 ± 0.23<sup>a</sup> | 1.18 ± 0.22<sup>a</sup> | 0.98 ± 0.42<sup>a</sup> | 2.41 ± 1.31<sup>a</sup> | 0.44 ± 0.12<sup>a</sup> |
| Spirosomaceae | 1.12 ± 0.19<sup>a</sup> | 1.34 ± 0.15<sup>a</sup> | 1.08 ± 0.32<sup>a</sup> | 0.76 ± 0.20<sup>a</sup> | 0.74 ± 0.22<sup>a</sup> |
| Pirellulaceae* | 1.27 ± 0.47<sup>a</sup> | 1.98 ± 0.41<sup>a</sup> | 0.61 ± 0.23<sup>a</sup> | 0.95 ± 0.25<sup>a</sup> | 0.11 ± 0.04<sup>a</sup> |
| Caulobacteraceae | 0.76 ± 0.09<sup>a</sup> | 1.36 ± 0.28<sup>a</sup> | 0.53 ± 0.03<sup>a</sup> | 0.73 ± 0.20<sup>a</sup> | 1.49 ± 0.64<sup>a</sup> |
| Other | 9.78 | 15.8 | 8.41 | 12.45 | 5.66 |

**Table 5:**
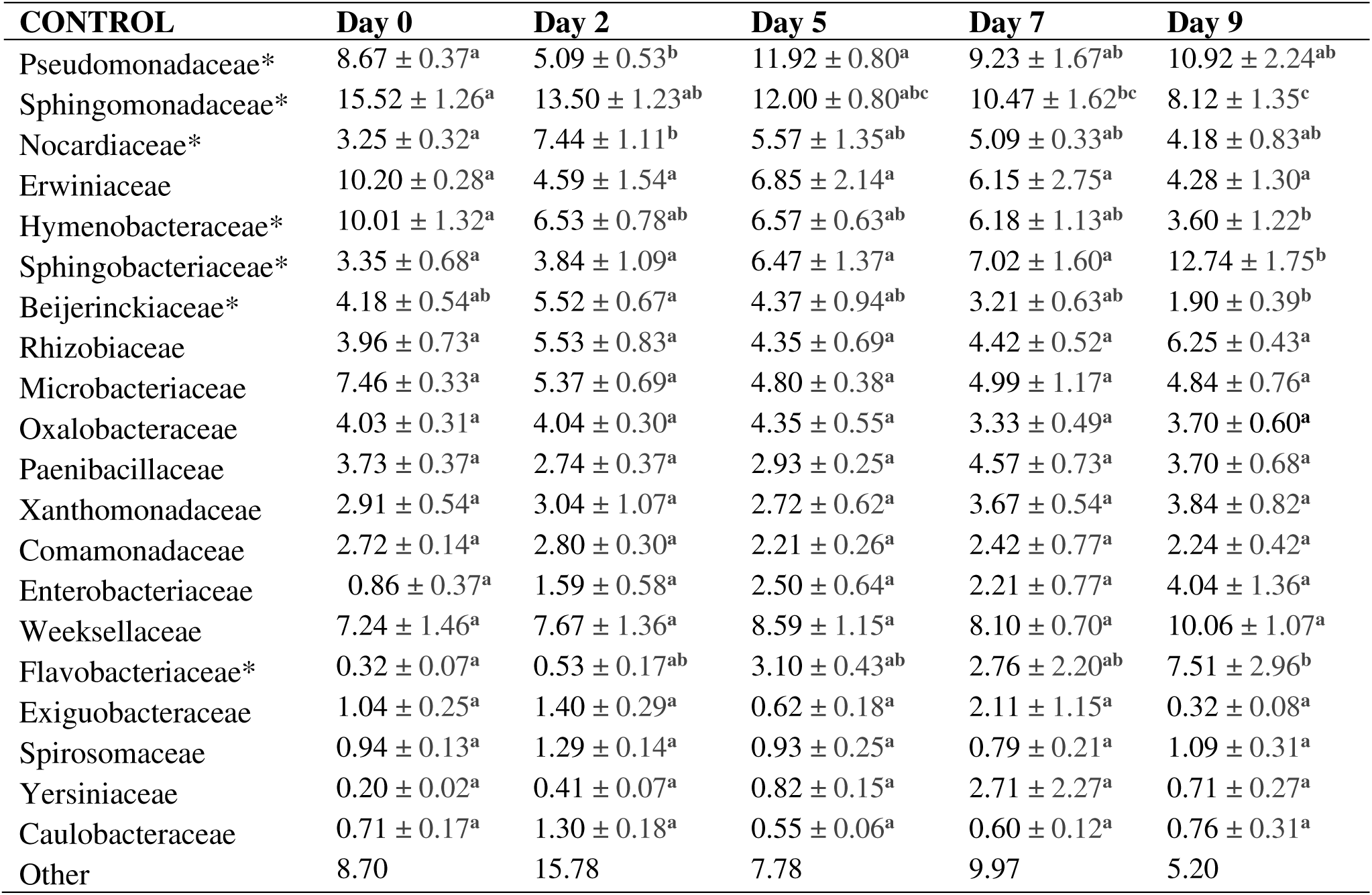
Average relative abundance (% ± standard error) of the 20 most abundant families (16S rDNA) of the CONTROL group across days 0, 2, 5. 7, and 9 (chloroplasts filtered with Qiime2, rarefied). * indicates an overall significant result was identified for that particular family. Letters a – e indicate significant differences over time. 17 common families between COMPETE, BLOCK and CONTROL groups are displayed first. All remaining lower abundant families are combined in “Other”.

| CONTROL | Day 0 | Day 2 | Day 5 | Day 7 | Day 9 |
| --- | --- | --- | --- | --- | --- |
| Pseudomonadaceae* | 8.67 ± 0.37<sup>a</sup> | 5.09 ± 0.53<sup>b</sup> | 11.92 ± 0.80<sup>a</sup> | 9.23 ± 1.67<sup>ab</sup> | 10.92 ± 2.24<sup>ab</sup> |
| Sphingomonadaceae* | 15.52 ± 1.26<sup>a</sup> | 13.50 ± 1.23<sup>ab</sup> | 12.00 ± 0.80<sup>abc</sup> | 10.47 ± 1.62<sup>bc</sup> | 8.12 ± 1.35<sup>c</sup> |
| Nocardiaceae* | 3.25 ± 0.32<sup>a</sup> | 7.44 ± 1.11<sup>b</sup> | 5.57 ± 1.35<sup>ab</sup> | 5.09 ± 0.33<sup>ab</sup> | 4.18 ± 0.83<sup>ab</sup> |
| Erwiniaceae | 10.20 ± 0.28<sup>a</sup> | 4.59 ± 1.54<sup>a</sup> | 6.85 ± 2.14<sup>a</sup> | 6.15 ± 2.75<sup>a</sup> | 4.28 ± 1.30<sup>a</sup> |
| Hymenobacteraceae* | 10.01 ± 1.32<sup>a</sup> | 6.53 ± 0.78<sup>ab</sup> | 6.57 ± 0.63<sup>ab</sup> | 6.18 ± 1.13<sup>ab</sup> | 3.60 ± 1.22<sup>b</sup> |
| Sphingobacteriaceae* | 3.35 ± 0.68<sup>a</sup> | 3.84 ± 1.09<sup>a</sup> | 6.47 ± 1.37<sup>a</sup> | 7.02 ± 1.60<sup>a</sup> | 12.74 ± 1.75<sup>b</sup> |
| Beijerinckiaceae* | 4.18 ± 0.54<sup>ab</sup> | 5.52 ± 0.67<sup>a</sup> | 4.37 ± 0.94<sup>ab</sup> | 3.21 ± 0.63<sup>ab</sup> | 1.90 ± 0.39<sup>b</sup> |
| Rhizobiaceae | 3.96 ± 0.73<sup>a</sup> | 5.53 ± 0.83<sup>a</sup> | 4.35 ± 0.69<sup>a</sup> | 4.42 ± 0.52<sup>a</sup> | 6.25 ± 0.43<sup>a</sup> |
| Microbacteriaceae | 7.46 ± 0.33<sup>a</sup> | 5.37 ± 0.69<sup>a</sup> | 4.80 ± 0.38<sup>a</sup> | 4.99 ± 1.17<sup>a</sup> | 4.84 ± 0.76<sup>a</sup> |
| Oxalobacteraceae | 4.03 ± 0.31<sup>a</sup> | 4.04 ± 0.30<sup>a</sup> | 4.35 ± 0.55<sup>a</sup> | 3.33 ± 0.49<sup>a</sup> | 3.70 ± 0.60<sup>a</sup> |
| Paenibacillaceae | 3.73 ± 0.37<sup>a</sup> | 2.74 ± 0.37<sup>a</sup> | 2.93 ± 0.25<sup>a</sup> | 4.57 ± 0.73<sup>a</sup> | 3.70 ± 0.68<sup>a</sup> |
| Xanthomonadaceae | 2.91 ± 0.54<sup>a</sup> | 3.04 ± 1.07<sup>a</sup> | 2.72 ± 0.62<sup>a</sup> | 3.67 ± 0.54<sup>a</sup> | 3.84 ± 0.82<sup>a</sup> |
| Comamonadaceae | 2.72 ± 0.14<sup>a</sup> | 2.80 ± 0.30<sup>a</sup> | 2.21 ± 0.26<sup>a</sup> | 2.42 ± 0.77<sup>a</sup> | 2.24 ± 0.42<sup>a</sup> |
| Enterobacteriaceae | 0.86 ± 0.37<sup>a</sup> | 1.59 ± 0.58<sup>a</sup> | 2.50 ± 0.64<sup>a</sup> | 2.21 ± 0.77<sup>a</sup> | 4.04 ± 1.36<sup>a</sup> |
| Weeksellaceae | 7.24 ± 1.46<sup>a</sup> | 7.67 ± 1.36<sup>a</sup> | 8.59 ± 1.15<sup>a</sup> | 8.10 ± 0.70<sup>a</sup> | 10.06 ± 1.07<sup>a</sup> |
| Flavobacteriaceae* | 0.32 ± 0.07<sup>a</sup> | 0.53 ± 0.17<sup>ab</sup> | 3.10 ± 0.43<sup>ab</sup> | 2.76 ± 2.20<sup>ab</sup> | 7.51 ± 2.96<sup>b</sup> |
| Exiguobacteraceae | 1.04 ± 0.25<sup>a</sup> | 1.40 ± 0.29<sup>a</sup> | 0.62 ± 0.18<sup>a</sup> | 2.11 ± 1.15<sup>a</sup> | 0.32 ± 0.08<sup>a</sup> |
| Spirosomaceae | 0.94 ± 0.13<sup>a</sup> | 1.29 ± 0.14<sup>a</sup> | 0.93 ± 0.25<sup>a</sup> | 0.79 ± 0.21<sup>a</sup> | 1.09 ± 0.31<sup>a</sup> |
| Yersiniaceae | 0.20 ± 0.02<sup>a</sup> | 0.41 ± 0.07<sup>a</sup> | 0.82 ± 0.15<sup>a</sup> | 2.71 ± 2.27<sup>a</sup> | 0.71 ± 0.27<sup>a</sup> |
| Caulobacteraceae | 0.71 ± 0.17<sup>a</sup> | 1.30 ± 0.18<sup>a</sup> | 0.55 ± 0.06<sup>a</sup> | 0.60 ± 0.12<sup>a</sup> | 0.76 ± 0.31<sup>a</sup> |
| Other | 8.70 | 15.78 | 7.78 | 9.97 | 5.20 |

### 3.6 Differentially abundant phyla and families

While the four major phyla (Proteobacteria, Bacteroidota, Actinobacteriota, and Firmicutes) made up over 90% of the bacterial communities, ANCOM identified 11 phyla from the remaining communities which were differentially abundant between the three groups i.e., Myxococcota, Patescibacteria, Cyanobacteria, Planctomycetota, Verrucomicrobiota, Acidobacteriota, Sumerlaeota, Bdellovibrionota, Deinococcota, Abditibacteriota, and Armatimonadota **(Table 6, top)**. While they were differentially abundant, their relative abundances were all well below 1% of average relative abundance with the exception of the Planctomycetota with a relative abundance of 2.20 % in BLOCK and 1.45 % in CONTROL but almost completely absent in COMPETE (0.03 %). The relative abundances of the differentially abundant phyla were closer for BLOCK and CONTROL than COMPETE. However, this observation is clearer at family level where ANCOM identified 19 differentially abundant families between all three groups **(Table 6, bottom)**. Again, all differentially abundant families have a relative abundance less than 1.25 %, except for Weeksellaceae which is one of the most abundant families **(Table 2)** of all three groups, but of lesser relative abundance in COMPETE (2.49 %) compared to BLOCK and CONTROL (8.72 and 8.33 %). Five of the identified families i.e., Pirellulaceae, Rubinisphaeraceae, Gemmataceae, Isosphaeraceae and uncultured Planctomycetales belong to the phylum Planctomycetota. Each of which were not present in the COMPETE group whereas they were present and had similar relative abundance levels in the BLOCK and CONTROL groups. Moreover, three families i.e., A4b, AKYG1722 and Caldilineaceae, belong to the phylum ‘Chloroflexi’ (Chloroflexota). These three families were also completely absent from COMPETE, yet present with similar relative abundances in the BLOCK and CONTROL groups.

**Table 6:** Average relative abundance (%) of the differentially abundant phyla and families identified by ANCOM (chloroplasts filtered with Qiime2, rarefied).

| **Phyla** | **COMPETE** | **BLOCK** | **CONTROL** |
| --- | --- | --- | --- |
| Myxococcota | 0.39 | 0.14 | 0.37 |
| Patescibacteria | 1.41 | 0.22 | 0.33 |
| Cyanobacteria | 0.00 | 0.25 | 0.10 |
| Planctomycetota | 0.03 | 2.20 | 1.45 |
| Verrucomicrobiota | 0.17 | 0.70 | 1.05 |
| Acidobacteriota | 0.08 | 0.63 | 0.47 |
| Sumerlaeota | 0.00 | 0.04 | 0.02 |
| Bdellovibrionota | 0.06 | 0.02 | 0.02 |
| Deinococcota | 1.22 | 0.65 | 0.43 |
| Abditibacteriota | 0.02 | 0.00 | 0.01 |
| Armatimonadota | 0.00 | 0.01 | 0.01 |
| **Families** | **COMPETE** | **BLOCK** | **CONTROL** |
| Pirellulaceae | 0.00 | 0.98 | 0.64 |
| Rubinisphaeraceae | 0.00 | 0.21 | 0.16 |
| Gemmataceae | 0.00 | 0.43 | 0.30 |
| Phormidiaceae | 0.00 | 0.19 | 0.07 |
| Isosphaeraceae | 0.00 | 0.22 | 0.11 |
| Planctomycetales; uncultured | 0.00 | 0.14 | 0.08 |
| Chitinophagaceae | 0.00 | 0.13 | 0.09 |
| Vicinamibacterales; uncultured | 0.02 | 0.28 | 0.21 |
| Saccharimonadaceae | 1.25 | 0.17 | 0.28 |
| Blastocatellaceae | 0.01 | 0.12 | 0.10 |
| Chthoniobacteraceae | 0.07 | 0.39 | 0.56 |
| A4b | 0.00 | 0.05 | 0.04 |
| Myxococcaceae | 0.37 | 0.11 | 0.32 |
| Xiphinematobacteraceae | 0.08 | 0.24 | 0.39 |
| Weeksellaceae | 2.49 | 8.72 | 8.33 |
| AKYG1722 | 0.00 | 0.03 | 0.03 |
| Mycobacteriaceae | 0.10 | 0.01 | 0.02 |
| Caldilineaceae | 0.00 | 0.16 | 0.16 |
| Solirubrobacteraceae | 0.05 | 0.01 | 0.02 |

Presence of the family Listeriaceae, which contain the genus *Listeria* was not detected in the majority of samples, even at the utilized sequence depth. A sequence depth of over 16,000 reads after rarefication should be able to identify at least 0.1 % relative abundant taxa, which would be the case for *L. monocytogenes*, when cultivation based abundances are taken into account (selective for *L. monocytogenes* as well as total heterotrophic counts (Culliney and Schmalenberger, 2022)). Consequently, a direct relationship of relative sequence abundances of certain phylogenetic groups over time alongside *L. monocytogenes* could not be analysed. Nevertheless, a qPCR approach confirmed the presence of *L. monocytogenes* in the utilized DNA samples, albeit at lower total quantities than determined via cultivation (data no shown).

## Discussion

The purpose of this study was to i) compare effectiveness of methods in reducing co-amplification of chloroplast DNA during 16S gene fragment amplification of prokaryotic DNA of a *L. monocytogenes* inoculated spinach phyllosphere, ii) report on their subsequent influence on bacterial community structures, iii) describe the progression of spinach’s phyllosphere community over time and iv) relate chloroplast DNA co-extraction and the associated phyllosphere community with growth of *L. monocytogenes*.

The present study on the spinach phyllosphere has demonstrated highly effective chloroplast blocking techniques. However, previous qPCR-based methods utilising universal primers 534F and 783R have proven effective at amplifying bacterial and mitochondrial DNA while limiting the amplification of plastid DNA (Rastogi et al., 2010). In that study, those primers enabled the accurate determination of total bacterial abundance of 10^5^ and 10^6^ g^−1^ of leaf of field-grown lettuce. The same qPCR protocol enumerated a total bacterial abundance of 5.1 to 7.7 log cfu g^-1^ of romaine lettuce (Ibekwe et al., 2013). While primer 783R is discriminating against chloroplast DNA through two mismatches, a significant co-amplification may still occur as identified by McManamon and colleagues ((McManamon et al., 2019), oral communication), thus further methods of chloroplast exclusions may be needed when the diversity of a community is to be analysed. Moreover, those authors revealed that determination of the cultivable total bacterial abundances using tryptic soy agar resulted in abundances of only 1 to 10% compared to qPCR estimates. Although, qPCR estimates may include non-viable bacterial cells recovered from the plant surface. The 783r primer has been be used in conjunction with the standard restriction fragment length polymorphism method for community analysis of spinach roots (Sakai et al., 2004). More recently, using the same two primers, qPCR estimates from two separate experiments of total bacteria of baby spinach’s phyllosphere were on average 7.1 and 6.4 log cfu g^-1^ (Truchado et al., 2019). It is noteworthy though that many common bacteria have multiple copies of 16S and therefore, a single bacterial cell can have 10 or even more copies (Schmalenberger et al., 2001) and therefore potentially inflate qPCR values. In contrast, by using cultivation-dependent techniques, TBC of spinach taken from the same sample asthe present study were similar to previously recorded qPCR abundances at day 0 around 7 log cfu g^-1^, increasing to approximately 8 log cfu g^-1^ thereafter (Culliney and Schmalenberger, 2022).

In this study, the pPNA clamp (BLOCK) was used in conjunction with primers covering the V1-V9 hypervariable region in the first PCR (outer primers), while the nested PCR then amplified the V3V4 region (inner primers) for Illumina sequencing. Other studies have employed the pPNA clamp only amplifying the V4 hypervariable region. The present study intentionally avoided this strategy in order to avoid potential PCR bias when the primers hybridizing with the target DNA in the nested PCR reaction are indeed in the primer locations of the first PCR or located at the position of the pPNA clamp.

Other studies have previously employed pPNA clamps for blocking of plastid DNA amplification for determination of food associated microbiomes. The use of a pPNA clamp with the amplification of the V4 region of bacterial 16S ribosomal RNA of rice seeds (*Oryza species* and *O. sativa* cultivars) avoided up losses of over 90% of sequences to chloroplasts and varied greatly not only at species level but also at cultivar level (Kim et al., 2020). Although chloroplast content for leafy vegetables are often similar, the pPNA method has also been used on other leafy type produce with larger chloroplast content e.g., fresh red alder leaves (Jackrel et al., 2017). A study that did not utilise a pPNA clamp retained a total of 77.4% and 11.6% chloroplast reads which were reduced to 4.84% and 0.21% with the inclusion of a pPNA clamp for terrestrial fresh red alder leaves and aquatic leaves, respectively. In the present study, the use of the same pPNA clamp (BLOCK) was highly effective in reducing chloroplast content from 10.93 to 0.26 % (40-fold). An additional study assessing the phyllosphere of tomato and strawberry leaves revealed the total percentage of non-bacterial reads were 96.6% (Legein et al., 2022). Utilising together pPNA and mPNA clamps reduced the percentage of non-bacterial reads to 39.6% while using pPNA alone only reduced the percentage to 81%. In the present study, COMPETE method was most successful in reducing chloroplast co-amplification (180-fold) compared to 40-fold for BLOCK. The RInvT primer (COMPETE) applied in the present study was developed for amplification of 16S rRNA gene fragments from lettuce bacterial communities for DGGE to avoid up to 90% chloroplast co-amplification as identified by direct amplicon sequencing in the same study (McManamon et al., 2019). However, a lettuce chloroplast DNA band was still visible after blocking in the fingerprint. Thus, utilisation of COMPETE was more effective with illumina sequencing of spinach bacterial communities. Therefore, pPNA (BLOCK) and COMPETE have shown to be less effective in situations where chloroplast content of produce has been higher than that of spinach (10.93 %) in the present study. The effectiveness of these methods could potentially reduce with increasing chloroplast content of spinach or differ for other similar produce with higher chloroplast content. Indeed, another study revealed similar chloroplast content of the spinach phyllo-epiphytic bacterial communities i.e., 3 to 15 % (Lopez-Velasco et al., 2011). However, cultivation factors such as nutrient solution fertigation and treatments with increasing nitrogen content (0 to 0.133 g nitrogen per kg soil) are positively correlated with both yield and chlorophyll content of spinach (Petropoulos et al., 2021, Xue and Yang, 2009). Chloroplast content of thicker (280 µM) sun exposed spinach leaves increased and remained higher than that of the thinner (186 µM) shaded leaves where chloroplast numbers reduced with depth (Cui et al., 1991). Chlorophyll content of leafy produce provides an indirect indication of the associated chloroplast concentration (Gedi et al., 2017).. When employing a pPNA clamp for certain plant microbiome studies, this effectiveness can be dramatically reduced owing to a single nucleotide mismatch between plastid DNA and the universal pPNA clamp used therefore, reducing deep amplicon sequencing (Fitzpatrick et al., 2018). Even when no mismatch occurs between plastid and host sequence, a high plastid content of 41% can still be observed on occasions as for instance reported for *Solanum dulcamara*, thus, indicating ineffectiveness of pPNA clamps for certain plant species regardless of host-pPNA match or mismatch (Fitzpatrick et al., 2018).

A promising alternative protocol incorporating a Ca9 nuclease and specific guide RNA (gRNA) to cut 16S rRNA targets during library construction removes host contamination from 61.2 to 70% to 4.5 to 11.3% during analysis of rice phyllosphere (Song and Xie, 2020). Although less chloroplast was present in this study, both methods (COMPETE and BLOCK) were more effective. The majority of studies mentioned in the present discussion which have targeted the spinach phyllosphere use bioinformatic software plugins for removal of chloroplast sequences post sequencing. These plugins were effective as also demonstrated in the present study, although their use is potentially resulting in large amounts of data loss when chloroplast co-amplification is high. Therefore, QIIME2 filtering of chloroplast could be applied independently without any prior blocking/exclusion effectively and without extra cost of blocking primers or pPNA clamps when co-extraction of chloroplast DNA is expected to be limited.

While low bacterial read numbers were no an issue in the current study, the use of pPNA clamps by Legein and colleagues (2022) did not resolve the issue of low bacterial read numbers i.e., 4,394 and 1,282 for strawberry and tomato leaf phyllospheres, respectively. The same authors revealed that bacterial densities were low at approximately 2 to 3 logs cfu g^-1^. Thus, suggesting PNA clamps may not be an appropriate method for other leafy vegetables expecting lower reads due to lower bacterial densities such as kale which started at approximately 2 to 3 logs compared to 7 to 8 logs for spinach (Culliney and Schmalenberger, 2022). However, similar to the observations of Kim and colleagues (2020), utilisation of PNA clamps did increase recovery of usable bacterial sequences by 5,395 reads belonging to the apple flower phytobiome (Steven et al., 2018). In contrast to Legein and colleagues (2022), nearly all non-bacterial reads belonged to plastid DNA whereas very little were mitochondria. Moreover, the most abundant bacterial OTUs were present in both the PNA clamp and control samples, suggesting utilisation of PNA clamps did not result in bias in reads recovered. In contrast, as a result of binding to the chloroplast blocking pPNA clamp, a strong sequencing bias against Proteobacterial 16S rRNA has previously been identified (Jackrel et al., 2017). It was suggested this occurred because of a conserved 14 base pair sequence in the Proteobacteria which matched the 17 base pair chloroplast blocking clamp. Those authors also provide a reference list of a further 1,500 taxa which contain that same 14 base pair sequence. However, such a bias against Proteobacterial 16S rRNA was not observed in the present study or by another study assessing pPNA clamp efficacy on 32 plant species (Fitzpatrick et al., 2018).

For most families present in the bacterial communities of the three presented groups, whether differentially abundant or not, similar relative abundances were identified. However, BLOCK and CONTROL were often most similar, except for the amount of chloroplast present prior to filtering. This suggests that the groups initially targeting the V1-V9 region had more similarities in terms of their complete 16S rRNA bacterial communities compared to the protocol that initially targeted the V2-V4 region, prior to adding Illumina adaptors. Based on this study, utilisation of the COMPETE strategy to describe the spinach phyllosphere has created a bias against amplification of the Plancomycetota phylum. Similarly, another study comparing microbial communities of Arctic environments using two different 16S rRNA primers revealed that the V4-V5 hypervariable region contained one third more ASVs belonging to Physcisphaerae and Plactomycetes classes, both of which belong to the Planctomycetota phylum, compared to the V3-V4 hypervariable region (Fadeev et al., 2021). Furthermore, the 783R primer which is used to amplify the V3-V4 hypervariable region is known to carry mismatches for targeting the phyla: Cyanobacteria, Planctomycetes, Chloroflexi, and Verrucomicrobia (Rastogi et al., 2010). This bias also extends towards chloroplast sequences which usually results in two mismatches and thus is often used intentionally to reduce chloroplast co-amplification. Interestingly, primers 806R and 783R greatly overlap (diverging names due to different numbering concepts i.e. counting from 3’ or 5’ end) with the former used for Illumina sequencing of the V3V4 region. However, both primers contain different ambiguities, hence the bias utilized with 783R is not identical to 806R.

In the present study, Chloroflexi and Verrucomicrobiota were underrepresented in the COMPETE group compared to both BLOCK and CONTROL groups, although, to a much lesser extent than the bias for Planctomycetota. Moreover, AKYG1722, A4b and Caldilineaceae families belonging to the Chloroflexi phylum were identified as being differentially abundant between all three tested amplification methods but were only completely absent from the COMPETE group. Chthoniobacteraceae and Xiphinematobacteraceae families belonging to Verrucomicrobiota were identified as being differentially abundant between all three groups, with relative abundance of both being least in the COMPETE group (0.10 %). In contrast, Saccharimonadaceae, Mycobacteriaceae and Solirubrobacteraceae were amplified by the RInvT primer (COMPETE) but to a lesser degree by BLOCK or CONTROL. Mycobacteriaceae and Solirubrobacteraceae biases may be of lesser importance due to their lower relative abundance (0.1 vs. 0.01%). While initial primer selection may have caused a certain level of bias, in the 2nd PCR we used the V3V4 primers for addition of illumina adaptors in all three approaches. Biases have been associated with nested PCR approaches compared to standard PCR for interpreting microbial community structures with Illumina MiSeq based 16S rRNA sequencing (Ercolini et al., 2015). However, these biases if present were dependent on the diversity in the samples and the number of cycles applied.

Studies have amplified different hypervariable regions for assessment of the spinach phyllosphere. V4 has been targeted using primer versions of 515F and 806R (Gu et al., 2020, Gu et al., 2018, Ibekwe et al., 2021). Other studies have used alternative primers for the same V4 hypervariable region such as a combination of three Roche 454 primers (Lopez-Velasco et al., 2013, Lopez-Velasco et al., 2011). Products of the resulting three amplifications were then pooled together in equal concentrations for amplicon sequencing. Tenzin and colleagues (2013) amplified the V3-V4 hypervariable region to describe the spinach phyllosphere using primers utilised by Klindworth and colleagues (2012) (Klindworth et al., 2013, Tenzin et al., 2020). However, the same region has been amplified with primers S-D-Bact-0341-b-S-17 and S-D-Bact-0785-a-A-21 with a second PCR to attach Illumina sequencing adaptors (Truchado et al., 2018), similar to the present study. Less common hypervariable regions amplified are covering the V5V6 region with primers 799F and 1115R which attempt to exclude cyanobacteria and therefore also exlude chloroplast co-amplification (Darlison et al., 2019). However, McManamon and colleagues found the exclusion of chloroplast amplicons insufficient without the addition of the COMPETE primer (McManamon et al., 2019). Another primer selection targeting the V6V8 region has been reported as well using primers B969F and BA1406R Targeting the V3/V4 or V4/V5 hypervariable regions provides high classification accuracies of 16S rRNA bacterial communities for next generation sequencing platforms (Claesson et al., 2010). However, the latter has been found to be better suited to minimise multiple different 16S rRNA gene sequences from a single type of organism, thus avoiding overestimation of alpha diversities (Schmalenberger et al., 2001). Nevertheless, the V3V4 target has been widely used in practise despite the reported bias potentials and will therefore likely remain the most popular choice from amplicons sequencing projects in the future.

As observed with their average relative abundances, BLOCK and CONTROL did not display any significant separations between groups across time while COMPETE had significant separations from both methods. In any case, all methods observed similar trends regarding alpha and beta diversity over time. Lopez-Velasco (2013) revealed after pyrosequencing the 16S rRNA gene amplicons (V4 region) of spinach that the alpha diversity metrics i.e., Shannon and evenness values (3.15 and 0.46, respectively) were lower than the present study (Lopez-Velasco et al., 2013). The lower diversity of their study is complimented with a 97% relative abundance of the Proteobacteria phylum which only ranged from approximately 50 to 60% in the current study. In their study, the relative abundance (%) of the other three most abundant phyla of this study i.e., Bacteroidota, Actinobacteriota and Firmicutes were either not reported or below 1% of relative abundance. In a separate study, the same authors used pyrosequencing to describe reductions in the diversity, richness, and evenness after packaging of the fresh spinach phyllosphere which were also lower than the present study i.e., their Shannon indexes ranged from 3.51 to 4.21 (Lopez-Velasco et al., 2011). Although, they noted an increase from storage at day 1 to day 15 at 4 °C. Albeit at a different storage profile, the present study had consistently the highest Shannon indexes at day 2 exceeding 6.1 independent of chosen method. By removing chloroplast DNA from NGS analysis of iceberg lettuce that made up to 90% of the sequence reads, the highest Shannon index of 2.4 increased to 4.3 (McManamon et al., 2019). However, in the present study, removal of chloroplast did not have such a strong influence on diversity, or richness as chloroplast co-amplification was comparatively lower. McManamon and colleagues (2019) noted after 7 days of storage, the composition of iceberg lettuce’s bacterial communities was considerably changed. These changes were also noted in the present study but earlier i.e., by day 5. Moreover, Lopez-Velasco and colleagues (2011) revealed that the fresh spinach phyllosphere had more similarity to the phyllosphere of packaged and stored produce at 4 °C after 1 and 15 days, compared to produce stored at 10 °C for 15 days. In another study, after 7 days storage of spinach leaves’ beta diversity (weighted UniFrac distances) had separated from day 0 and bacterial communities of 10 and 15 °C were more similar than 4 °C (Gu et al., 2018). Those studies reveal that increases in time and storage temperature both influenced bacterial community structures with temperatures beyond refrigeration temperatures causing greater separations in beta diversity. In the present study, there was an increase in storage temperature from 7 to 10 °C from days 6 to 9. Therefore, both time and storage temperature may be responsible for changes in the bacterial community across time. Indeed, cluster analysis revealed that bacterial communities of spinach were only 50 to 60 % similar at temperatures profiles of 4 and 10 °C (Lopez-Velasco et al., 2010). Moreover at both temperatures separately, DGGE patterns indicated changes throughout storage occur as days 0 and 5 were more similar to each other than days 10 and 15 (Lopez-Velasco et al., 2010).

Additionally, the change in the structure of the spinach phyllosphere over time is important to fully understand the behaviour of *L. monocytogenes* on leafy vegetables. From day 7 to 9, chloroplast content decreased to 2.64 % while the associated *L. monocytogenes’* populations increased drastically (Culliney and Schmalenberger, 2022). Thus, while blocking methods were effective, chloroplast content also naturally reduced throughout storage to low contents by day 9. Decreases in chloroplast content can result from continued cell expansion without plastid DNA replication or due to degradation of chloroplasts (Buet et al., 2019, Tymms et al., 1983). The proportion of chloroplast to total DNA content i.e., 19.6 to 23.4 % of expanding spinach leaves remained constant with increasing leaf size from 2 to 10 cm (Scott and Possingham, 1980). Conversely, when spinach leaves were detached and aged over 4 days, leaf chlorophyll content reduced by 20 % and by a further 30 % when supplied with glucose (Kilb et al., 1996). Other studies have also demonstrated a reduction of plastids of detached spinach leaves throughout storage (Wrischer, 1978). However, this occurred more strongly under conditions of darkness compared to light. Similarly, senescence induced by shading reduced chloroplast content to total DNA content from 19 to 15% and nitrogen deficiency in tandem with yellowing of leaf and senescence reduced chloroplast to total DNA content from 14.1 to 8.5% after 11 days (Scott and Possingham, 1983).

The present study was able to identify the presence of bacteria, and their shifts in abundance, important to the growth of *L. monocytogenes*. For example, bacteria from the family Pseudomonadaceae family which were of highly abundant (4.37 to 15.43 %), are associated with hydrolysis of proteins into amino acids which induce the stimulation of *L. monocytogenes* growth (Marshall et al., 1992). Contrariwise, Lactobacillales (order) that were not completely absent but were present in low abundance (0 to 1.19 %; data not shown in results), due to their competitive growth abilities are commonly associated with decreased *L. monocytogenes* survival (Østergaard et al., 2014). Indeed, the putative *L. monocytogenes* growth enhancing *Pseudomonas* bacteria have previously been associated with spinach leaves of neutral pH (Babic et al., 1996). Additionally, as *Pseudomonas* species are pectolytic their presence is positively correlated with the degradation and spoilage of such leafy vegetables which increases during storage. With an overall relative abundance of 35 to 53%, *Pseudomonas* has been referred to as the most commonly occurring genus of the spinach and rocket phyllospheres even after being harvested in different seasons (spring and autumn) (Rosberg et al., 2021). In the present study, while COMPETE had significantly higher relative abundance of Pseudomonoadaceae compared to BLOCK and CONTROL, all three groups demonstrated the same trends in Pseudomonoadaceae content. All three groups also displayed similar trends of increasing Lactobacillales content from day 0 (0.23 to 0.32 %) to 5 (0.76 to 1.19 %) followed by decreases until day 9 near absence. The largest decrease in Lactobacillales content from day 7 (0.49 to 0.64%) to 9 (0.03 to 0.04 %) was associated with the greatest increase in *L. monocytogenes* i.e., 1.10 log_10_ cfu g^-1^ (Culliney and Schmalenberger, 2022). Thus, the reduction of Lactobacillales (which are associated with decreasing *L. monocytogenes* growth) content to near absence in combination with an increase in Pseudomonodaceae content (+ 1.69 to 2.34 increase across three methods from day 7 to 9) may have been responsible for the greater increase in *L. monocytogenes* growth observed alongside the higher storage temperate from day6-9.

## Conclusion

In conclusion, this study provides evidence that both BLOCK and especially COMPETE were highly effective methods in reducing co-amplification of chloroplast DNA. Although, spinach’s chloroplast DNA content was relatively low (10.93 %) and number of bacterial reads were high. Therefore, utilisation of these methods may not be necessary for spinach in future studies, and filter-table methods in the QIIME2-taxa plugin will suffice as a chloroplast removing method. In the present study, primer selection was once more highlighted as an important factor in amplification bias. When using the COMPETE solution for exclusion of chloroplast co-amplification, this needs to be thoroughly considered. In contrast, the BLOCK solution while less effective in chloroplast co-amplification suppression may result in fewer amplicon biases due to the flexibility of choosing the fist primer set, which is not an option for the COMPETE solution. Researchers may utilise these effective approaches in future studies on leafy produce with higher chloroplast co-extraction or low phyllosphere microbial abundance.

The current study also revealed reductions in chloroplast co-extraction across time and presence of phyllosphere community members specifically Pseudomonadaceae and Lactobacillales, which all potentially influence growth of *L. monocytogenes* on leafy vegetables. Therefore, future studies should determine which is the causative agent or if both are responsible, where conflicting *L. monocytogenes* growth data is available for spinach and various other plant species i.e., rocket and kale produce from differing cultivation methods (Culliney and Schmalenberger, 2022). Moreover, further research needs to assess whether higher diversity of the phyllosphere reduces the competitiveness of invading species (Darlison et al., 2019), such as *L. monocytogenes*.

## Acknowledgements

We would like to thank the Department of Agriculture, Food and the Marine (DAFM) for funding this project under the consortium ListeriaChallengeStudies (grant number 17F/244).

## Bibliography

Anderson, M. J. & Walsh, D. C. I. 2013. Permanova, Anosim, and the Mantel test in the face of heterogeneous dispersions: What null hypothesis are you testing? Ecological Monographs, 83, 557–574.

Babic, I., Roy, S., Watada, A. E. & Wergin, W. P. 1996. Changes in microbial populations on fresh cut spinach. International Journal of Food Microbiology, 31, 107–119.

Bolyen, E., Rideout, J. R., Dillon, M. R., Bokulich, N. A., Abnet, C. C., Al-Ghalith, G. A., Alexander, H., Alm, E. J., Arumugam, M., Asnicar, F., Bai, Y., Bisanz, J. E., Bittinger, K., Brejnrod, A., Brislawn, C. J., Brown, C. T., Callahan, B. J., Caraballo-Rodríguez, A. M., Chase, J., Cope, E. K., Da Silva, R., Diener, C., Dorrestein, P. C., Douglas, G. M., Durall, D. M., Duvallet, C., Edwardson, C. F., Ernst, M., Estaki, M., Fouquier, J., Gauglitz, J. M., Gibbons, S. M., Gibson, D. L., Gonzalez, A., Gorlick, K., Guo, J., Hillmann, B., Holmes, S., Holste, H., Huttenhower, C., Huttley, G. A., Janssen, S., Jarmusch, A. K., Jiang, L., Kaehler, B. D., Kang, K. B., Keefe, C. R., Keim, P., Kelley, S. T., Knights, D., Koester, I., Kosciolek, T., Kreps, J., Langille, M. G. I., Lee, J., Ley, R., Liu, Y.-X., Loftfield, E., Lozupone, C., Maher, M., Marotz, C., Martin, B. D., Mcdonald, D., Mciver, L. J., Melnik, A. V., Metcalf, J. L., Morgan, S. C., Morton, J. T., Naimey, A. T., Navas-Molina, J. A., Nothias, L. F., Orchanian, S. B., Pearson, T., Peoples, S. L., Petras, D., Preuss, M. L., Pruesse, E., Rasmussen, L. B., Rivers, A., Robeson, M. S., Rosenthal, P., Segata, N., Shaffer, M., Shiffer, A., Sinha, R., Song, S. J., Spear, J. R., Swafford, A. D., Thompson, L. R., Torres, P. J., Trinh, P., Tripathi, A., Turnbaugh, P. J., Ul-Hasan, S., Van Der Hooft, J. J. J., Vargas, F., Vázquez-Baeza, Y., Vogtmann, E., Von Hippel, M., Walters, W., et al. 2019. Author Correction: Reproducible, interactive, scalable and extensible microbiome data science using Qiime 2. Nature Biotechnology, 37, 1091–1091.

Buet, A., Costa, M. L., Martínez, D. E. & Guiamet, J. J. 2019. Chloroplast Protein Degradation in Senescing Leaves: Proteases and Lytic Compartments. Frontiers in Plant Science, 10.

Claesson, M. J., Wang, Q., O’sullivan, O., Greene-Diniz, R., Cole, J. R., Ross, R. P. & O’toole, P. W. 2010. Comparison of two next-generation sequencing technologies for resolving highly complex microbiota composition using tandem variable 16s rrna gene regions. Nucleic Acids Research, 38, e200–e200.

Cui, M., Vogelmann, T. C. & Smith, W. K. 1991. Chlorophyll and light gradients in sun and shade leaves of Spinacia oleracea. Plant, Cell and Environment, 14, 493–500.

Culliney, P. & Schmalenberger, A. 2022. Cultivation Conditions of Spinach and Rocket Influence Epiphytic Growth of Listeria monocytogenes. Foods, 11.

Daniell, H., Lin, C.-S., Yu, M. & Chang, W.-J. 2016. Chloroplast genomes: diversity, evolution, and applications in genetic engineering. Genome Biology, 17.

Darlison, J., Mogren, L., Rosberg, A. K., Grudén, M., Minet, A., Liné, C., Mieli, M., Bengtsson, T., Håkansson, Å., Uhlig, E., Becher, P. G., Karlsson, M. & Alsanius, B. W. 2019. Leaf mineral content govern microbial community structure in the phyllosphere of spinach (Spinacia oleracea) and rocket (Diplotaxis tenuifolia). Science of The Total Environment, 675, 501–512.

Dastogeer, K. M. G., Tumpa, F. H., Sultana, A., Akter, M. A. & Chakraborty, A. 2020. Plant microbiome–an account of the factors that shape community composition and diversity. Current Plant Biology, 23.

Ercolini, D., Yu, G., Fadrosh, D., Goedert, J. J., Ravel, J. & Goldstein, A. M. 2015. Nested Pcr Biases in Interpreting Microbial Community Structure in 16s rrna Gene Sequence Datasets. Plos One, 10.

Eurl Lm 2019. Technical guidance document for conducting shelf-life studies on *Listeria monocytogenes* in ready-to-eat foods Version 3-Amended (21/02/2019). Maisons-Alfort, France: Eurl Listeria monocytogenes, Anses. .

Eurl Lm 2021. Technical guidance document for conducting shelf-life studies on *Listeria monocytogenes* in ready-to-eat foods Version 4-Amended (01/07/2021). Maisons-Alfort, France: Eurl *Listeria* monocytogenes, Anses. .

Fadeev, E., Cardozo-Mino, M. G., Rapp, J. Z., Bienhold, C., Salter, I., Salman-Carvalho, V., Molari, M., Tegetmeyer, H. E., Buttigieg, P. L. & Boetius, A. 2021. Comparison of Two 16s rrna Primers (V3–V4 and V4–V5) for Studies of Arctic Microbial Communities. Frontiers in Microbiology, 12.

Faith, D. P. 1992. Conservation evaluation and phylogenetic diversity. Biological Conservation, 61, 1–10.

Fitzpatrick, C. R., Lu-Irving, P., Copeland, J., Guttman, D. S., Wang, P. W., Baltrus, D. A., Dlugosch, K. M. & Johnson, M. T. J. 2018. Chloroplast sequence variation and the efficacy of peptide nucleic acids for blocking host amplification in plant microbiome studies. Microbiome, 6.

Francis, G. A. & O’beirne, D. 2002. Effects of the indigenous microflora of minimally processed lettuce on the survival and growth of*Listeria innocua*. Int. J. Food Sci. Tech., 33, 477–488.

Gao, C., Ren, X., Mason, A. S., Liu, H., Xiao, M., Li, J. & Fu, D. 2013. Horizontal gene transfer in plants. Functional & Integrative Genomics, 14, 23–29.

Gedi, M. A., Briars, R., Yuseli, F., Zainol, N., Darwish, R., Salter, A. M. & Gray, D. A. 2017. Component analysis of nutritionally rich chloroplasts: recovery from conventional and unconventional green plant species. Journal of Food Science and Technology, 54, 2746–2757.

Gong, T. & Xin, X. F. 2021. Phyllosphere microbiota: Community dynamics and its interaction with plant hosts. Journal of Integrative Plant Biology, 63, 297–304.

Gu, G., Ottesen, A., Bolten, S., Luo, Y., Rideout, S. & Nou, X. 2020. Microbiome convergence following sanitizer treatment and identification of sanitizer resistant species from spinach and lettuce rinse water. International Journal of Food Microbiology, 318.

Gu, G., Ottesen, A., Bolten, S., Ramachandran, P., Reed, E., Rideout, S., Luo, Y., Patel, J., Brown, E. & Nou, X. 2018. Shifts in spinach microbial communities after chlorine washing and storage at compliant and abusive temperatures. Food Microbiology, 73, 73–84.

Hanshew, A. S., Mason, C. J., Raffa, K. F. & Currie, C. R. 2013. Minimization of chloroplast contamination in 16s rrna gene pyrosequencing of insect herbivore bacterial communities. Journal of Microbiological Methods, 95, 149–155.

Ibekwe, A. M., Ors, S., Ferreira, J. F. S., Liu, X. & Suarez, D. L. 2021. Influence of seasonal changes and salinity on spinach phyllosphere bacterial functional assemblage. Plos One, 16.

Ibekwe, A. M., Williams, T. R., Moyne, A.-L., Harris, L. J. & Marco, M. L. 2013. Season, Irrigation, Leaf Age, and Escherichia coli Inoculation Influence the Bacterial Diversity in the Lettuce Phyllosphere. PLos One, 8.

Jackrel, S. L., Owens, S. M., Gilbert, J. A. & Pfister, C. A. 2017. Identifying the plant-associated microbiome across aquatic and terrestrial environments: the effects of amplification method on taxa discovery. Molecular Ecology Resources, 17, 931–942.

Kilb, B., Wietoska, H. & Godde, D. 1996. Changes in the expression of photosynthetic genes precede loss of photosynthetic activities and chlorophyll when glucose is supplied to mature spinach leaves. Plant Science, 115, 225–235.

Kim, H., Lee, K. K., Jeon, J., Harris, W. A. & Lee, Y.-H. 2020. Domestication of Oryza species eco-evolutionarily shapes bacterial and fungal communities in rice seed. Microbiome, 8.

Klindworth, A., Pruesse, E., Schweer, T., Peplies, J., Quast, C., Horn, M. & Glöckner, F. O. 2013. Evaluation of general 16s ribosomal Rna gene Pcr primers for classical and next-generation sequencing-based diversity studies. Nucleic Acids Research, 41, e1–e1.

Knief, C. 2014. Analysis of plant microbe interactions in the era of next generation sequencing technologies. Frontiers in Plant Science, 5.

Legein, M., Smets, W., Wuyts, K., Bosmans, L., Samson, R., Lebeer, S. & Deangelis, K. M. 2022. The Greenhouse Phyllosphere Microbiome and Associations with Introduced Bumblebees and Predatory Mites. Microbiology Spectrum, 10.

Lindow, S. E. & Brandl, M. T. 2003. Microbiology of the Phyllosphere. Applied and Environmental Microbiology, 69, 1875–1883.

Lopez-Velasco, G., Carder, P. A., Welbaum, G. E. & Ponder, M. A. 2013. Diversity of the spinach (Spinacia oleracea) spermosphere and phyllosphere bacterial communities. Fems Microbiology Letters, 346, 146–154.

Lopez-Velasco, G., Davis, M., Boyer, R. R., Williams, R. C. & Ponder, M. A. 2010. Alterations of the phylloepiphytic bacterial community associated with interactions of Escherichia coli O157:H7 during storage of packaged spinach at refrigeration temperatures. Food Microbiology, 27, 476–486.

Lopez-Velasco, G., Welbaum, G. E., Boyer, R. R., Mane, S. P. & Ponder, M. A. 2011. Changes in spinach phylloepiphytic bacteria communities following minimal processing and refrigerated storage described using pyrosequencing of 16s rrna amplicons. Journal of Applied Microbiology, 110, 1203–1214.

Lozupone, C. & Knight, R. 2005. UniFrac: a New Phylogenetic Method for Comparing Microbial Communities. Applied and Environmental Microbiology, 71, 8228–8235.

Lozupone, C. A., Hamady, M., Kelley, S. T. & Knight, R. 2007. Quantitative and Qualitative β Diversity Measures Lead to Different Insights into Factors That Structure Microbial Communities. Applied and Environmental Microbiology, 73, 1576–1585.

Marshall, D. L., Andrews, L. S., Wells, J. H. & Farr, A. J. 1992. Influence of modified atmosphere packaging on the competitive growth of Listeria monocytogenes and Pseudomonas fluorescens on precooked chicken. Food Microbiology, 9, 303–309.

Mcmanamon, O., Kaupper, T., Scollard, J. & Schmalenberger, A. 2019. Nisin application delays growth of Listeria monocytogenes on fresh-cut iceberg lettuce in modified atmosphere packaging, while the bacterial community structure changes within one week of storage. Postharvest Biology and Technology, 147, 185–195.

Østergaard, N. B., Eklöw, A. & Dalgaard, P. 2014. Modelling the effect of lactic acid bacteria from starter- and aroma culture on growth of Listeria monocytogenes in cottage cheese. International Journal of Food Microbiology, 188, 15–25.

Pellestor, F. & Paulasova, P. 2004. The peptide nucleic acids (PNAs), powerful tools for molecular genetics and cytogenetics. European Journal of Human Genetics, 12, 694–700.

Petropoulos, S. A., El-Nakhel, C., Graziani, G., Kyriacou, M. C. & Rouphael, Y. 2021. The Effects of Nutrient Solution Feeding Regime on Yield, Mineral Profile, and Phytochemical Composition of Spinach Microgreens. Horticulturae, 7.

Rastogi, G., Coaker, G. L. & Leveau, J. H. J. 2013. New insights into the structure and function of phyllosphere microbiota through high-throughput molecular approaches. Fems Microbiology Letters, 348, 1–10.

Rastogi, G., Tech, J. J., Coaker, G. L. & Leveau, J. H. J. 2010. A Pcr-based toolbox for the culture-independent quantification of total bacterial abundances in plant environments. Journal of Microbiological Methods, 83, 127–132.

Rosberg, A. K., Darlison, J., Mogren, L. & Alsanius, B. W. 2021. Commercial wash of leafy vegetables do not significantly decrease bacterial load but leads to shifts in bacterial species composition. Food Microbiology, 94.

Sakai, M., Matsuka, A., Komura, T. & Kanazawa, S. 2004. Application of a new Pcr primer for terminal restriction fragment length polymorphism analysis of the bacterial communities in plant roots. Journal of Microbiological Methods, 59, 81–89.

Schmalenberger, A., Schwieger, F. & Tebbe, C. C. 2001. Effect of Primers Hybridizing to Different Evolutionarily Conserved Regions of the Small-Subunit rrna Gene in Pcr-Based Microbial Community Analyses and Genetic Profiling. Applied and Environmental Microbiology, 67, 3557–3563.

Scott, N. S. & Possingham, J. V. 1980. Chloroplast Dna in Expanding Spinach Leaves. Journal of Experimental Botany, 31, 1081–1092.

Scott, N. S. & Possingham, J. V. 1983. Changes in Chloroplast Dna Levels during Growth of Spinach Leaves. Journal of Experimental Botany, 34, 1756–1767.

Song, L. & Xie, K. 2020. Engineering Crispr/Cas9 to mitigate abundant host contamination for 16s rrna gene-based amplicon sequencing. Microbiome, 8.

Steven, B., Huntley, R. B. & Zeng, Q. 2018. The Influence of Flower Anatomy and Apple Cultivar on the Apple Flower Phytobiome. Phytobiomes Journal, 2, 171–179.

Stone, B. W. G., Weingarten, E. A. & Jackson, C. R. 2018. The Role of the Phyllosphere Microbiome in Plant Health and Function. Annual Plant Reviews online.

Tenzin, S., Ogunniyi, A. D., Ferro, S., Deo, P. & Trott, D. J. 2020. Effects of an Eco-Friendly Sanitizing Wash on Spinach Leaf Bacterial Community Structure and Diversity. Applied Sciences, 10.

Truchado, P., Gil, M. I., Moreno-Candel, M. & Allende, A. 2019. Impact of weather conditions, leaf age and irrigation water disinfection on the major epiphytic bacterial genera of baby spinach grown in an open field. Food Microbiology, 78, 46–52.

Truchado, P., Gil, M. I., Suslow, T. & Allende, A. 2018. Impact of chlorine dioxide disinfection of irrigation water on the epiphytic bacterial community of baby spinach and underlying soil. Plos One, 13.

Tymms, M. J., Scott, N. S. & Possingham, J. V. 1983. Dna Content of Beta vulgaris Chloroplasts during Leaf Cell Expansion. Plant Physiology, 71, 785–788.

Vorholt, J. A. 2012. Microbial life in the phyllosphere. Nature Reviews Microbiology, 10, 828–840.

Wrischer, M. 1978. Ultrastructural Changes in Plastids of Detached Spinach Leaves. Zeitschrift für Pflanzenphysiologie, 86, 95–106.

Xu, N., Zhao, Q., Zhang, Z., Zhang, Q., Wang, Y., Qin, G., Ke, M., Qiu, D., Peijnenburg, W. J. G. M., Lu, T. & Qian, H. 2022. Phyllosphere Microorganisms: Sources, Drivers, and Their Interactions with Plant Hosts. Journal of Agricultural and Food Chemistry, 70, 4860–4870.

Xue, L. & Yang, L. 2009. Deriving leaf chlorophyll content of green-leafy vegetables from hyperspectral reflectance. Isprs Journal of Photogrammetry and Remote Sensing, 64, 97–106.

