## Supplementary for "Bacterial community structure analysis on *Listeria monocytogenes* inoculated spinach leaves is affected by PCR based methods to exclude chloroplast co-amplification"

### Supplementary Information

**Supplementary Information: Table 1:** Dada2 run statistics containing number of input, filtered, % of input passed filter, denoised, merged, % of input merged, non-chimeric and % of input non-chimeric reads.

| sample-id | input | filtered | % of input passed filter | denoised | merged | % of input merged | non-chimeric | % of input non-chimeric |
|---|---|---|---|---|---|---|---|---|
| RInvTDay0Rep1 | 51,325 | 44,973 | 87.62 | 43,803 | 41,196 | 80.26 | 36,429 | 70.98 |
| RInvTDay0Rep2 | 74,334 | 65,654 | 88.32 | 64,537 | 61,142 | 82.25 | 53,079 | 71.41 |
| RInvTDay0Rep3 | 91,304 | 78,996 | 86.52 | 77,071 | 71,543 | 78.36 | 62,357 | 68.3 |
| RInvTDay0Rep4 | 81,262 | 70,446 | 86.69 | 68,463 | 63,511 | 78.16 | 55,768 | 68.63 |
| RInvTDay2Rep1 | 73,753 | 64,884 | 87.97 | 62,939 | 57,793 | 78.36 | 53,244 | 72.19 |
| RInvTDay2Rep2 | 76,738 | 67,650 | 88.16 | 65,881 | 60,701 | 79.1 | 54,705 | 71.29 |
| RInvTDay2Rep3 | 92,148 | 79,853 | 86.66 | 77,451 | 70,693 | 76.72 | 64,129 | 69.59 |
| RInvTDay2Rep4 | 113,629 | 101,694 | 89.5 | 99,428 | 92,359 | 81.28 | 81,455 | 71.69 |
| RInvTDay5Rep1 | 97,816 | 87,230 | 89.18 | 85,459 | 79,818 | 81.6 | 67,493 | 69 |
| RInvTDay5Rep2 | 84,184 | 73,219 | 86.97 | 71,389 | 66,193 | 78.63 | 57,477 | 68.28 |
| RInvTDay5Rep3 | 80,225 | 70,476 | 87.85 | 68,583 | 62,804 | 78.28 | 54,032 | 67.35 |
| RInvTDay5Rep4 | 92,117 | 81,127 | 88.07 | 79,680 | 74,963 | 81.38 | 66,715 | 72.42 |
| RInvTDay7Rep1 | 49,604 | 43,253 | 87.2 | 42,209 | 40,001 | 80.64 | 36,153 | 72.88 |
| RInvTDay7Rep2 | 88,136 | 78,183 | 88.71 | 76,701 | 71,736 | 81.39 | 63,494 | 72.04 |
| RInvTDay7Rep3 | 83,638 | 72,914 | 87.18 | 71,573 | 67,128 | 80.26 | 57,795 | 69.1 |
| RInvTDay7Rep4 | 81,765 | 70,819 | 86.61 | 69,370 | 64,759 | 79.2 | 58,362 | 71.38 |
| RInvTDay9Rep1 | 73,272 | 64,886 | 88.55 | 63,947 | 60,193 | 82.15 | 48,824 | 66.63 |
| RInvTDay9Rep2 | 66,781 | 58,632 | 87.8 | 57,385 | 53,468 | 80.06 | 47,343 | 70.89 |
| RInvTDay9Rep3 | 91,185 | 79,892 | 87.62 | 78,461 | 73,168 | 80.24 | 61,244 | 67.16 |
| RInvTDay9Rep4 | 83,691 | 74,663 | 89.21 | 73,429 | 67,793 | 81 | 57,882 | 69.16 |
| pPNADay0Rep1 | 82,442 | 72,560 | 88.01 | 70,708 | 65,394 | 79.32 | 57,032 | 69.18 |
| pPNADay0Rep2 | 73,030 | 64,023 | 87.67 | 62,188 | 57,256 | 78.4 | 50,575 | 69.25 |
| pPNADay0Rep3 | 85,736 | 74,870 | 87.33 | 72,756 | 67,570 | 78.81 | 61,982 | 72.29 |
| pPNADay0Rep4 | 108,341 | 92,193 | 85.1 | 90,422 | 85,095 | 78.54 | 77,277 | 71.33 |
| pPNADay2Rep1 | 68,899 | 57,717 | 83.77 | 56,134 | 50,820 | 73.76 | 48,240 | 70.02 |
| pPNADay2Rep2 | 113,473 | 99,919 | 88.06 | 97,045 | 87,461 | 77.08 | 77,894 | 68.65 |
| pPNADay2Rep3 | 81,585 | 69,983 | 85.78 | 67,674 | 60,545 | 74.21 | 56,959 | 69.82 |
| pPNADay2Rep4 | 97,022 | 85,049 | 87.66 | 82,963 | 76,000 | 78.33 | 69,792 | 71.93 |
| pPNADay5Rep1 | 90,399 | 36,127 | 39.96 | 35,137 | 29,900 | 33.08 | 27,323 | 30.22 |
| pPNADay5Rep2 | 64,722 | 56,435 | 87.2 | 54,501 | 50,003 | 77.26 | 45,487 | 70.28 |
| pPNADay5Rep3 | 106,488 | 93,158 | 87.48 | 91,043 | 84,415 | 79.27 | 75,398 | 70.8 |
| pPNADay5Rep4 | 108,226 | 89,751 | 82.93 | 88,176 | 82,081 | 75.84 | 73,942 | 68.32 |
| pPNADay7Rep1 | 99,757 | 84,411 | 84.62 | 82,881 | 77,056 | 77.24 | 69,415 | 69.58 |
| pPNADay7Rep2 | 69,432 | 60,506 | 87.14 | 59,170 | 54,987 | 79.2 | 50,601 | 72.88 |
| pPNADay7Rep3 | 81,844 | 72,326 | 88.37 | 70,253 | 64,883 | 79.28 | 54,278 | 66.32 |
| pPNADay7Rep4 | 59,276 | 46,490 | 78.43 | 45,312 | 41,390 | 69.83 | 38,549 | 65.03 |
| pPNADay9Rep1 | 45,076 | 36,916 | 81.9 | 36,247 | 34,566 | 76.68 | 31,997 | 70.98 |
| pPNADay9Rep2 | 48,777 | 37,757 | 77.41 | 36,598 | 33,486 | 68.65 | 31,381 | 64.34 |

| | | | | | | | | |
|---|---|---|---|---|---|---|---|---|
| pPNADay9Rep3 | 55,709 | 43,207 | 77.56 | 41,936 | 38,550 | 69.2 | 35,104 | 63.01 |
| pPNADay9Rep4 | 49,753 | 37,170 | 74.71 | 36,110 | 32,293 | 64.91 | 28,908 | 58.1 |
| ControlDay0Rep1 | 49,453 | 39,769 | 80.42 | 38,388 | 35,461 | 71.71 | 31,950 | 64.61 |
| ControlDay0Rep2 | 33,899 | 24,842 | 73.28 | 23,751 | 21,567 | 63.62 | 20,024 | 59.07 |
| ControlDay0Rep3 | 62,560 | 48,933 | 78.22 | 47,176 | 42,993 | 68.72 | 40,374 | 64.54 |
| ControlDay0Rep4 | 73,136 | 63,604 | 86.97 | 61,871 | 57,719 | 78.92 | 50,409 | 68.93 |
| ControlDay2Rep1 | 56,018 | 44,411 | 79.28 | 42,635 | 38,221 | 68.23 | 36,457 | 65.08 |
| ControlDay2Rep2 | 50,370 | 38,258 | 75.95 | 36,637 | 33,053 | 65.62 | 30,817 | 61.18 |
| ControlDay2Rep3 | 48,490 | 36,383 | 75.03 | 34,563 | 30,721 | 63.36 | 29,835 | 61.53 |
| ControlDay2Rep4 | 116,976 | 103,686 | 88.64 | 101,072 | 92,723 | 79.27 | 84,739 | 72.44 |
| ControlDay5Rep1 | 74,560 | 65,726 | 88.15 | 63,846 | 58,854 | 78.94 | 52,105 | 69.88 |
| ControlDay5Rep2 | 78,458 | 69,164 | 88.15 | 67,625 | 62,988 | 80.28 | 55,986 | 71.36 |
| ControlDay5Rep3 | 90,440 | 78,706 | 87.03 | 77,155 | 71,798 | 79.39 | 64,420 | 71.23 |
| ControlDay5Rep4 | 101,056 | 88,529 | 87.6 | 86,897 | 80,941 | 80.1 | 71,340 | 70.59 |
| ControlDay7Rep1 | 78,242 | 69,360 | 88.65 | 67,478 | 62,539 | 79.93 | 53,977 | 68.99 |
| ControlDay7Rep2 | 77,137 | 68,137 | 88.33 | 66,356 | 61,847 | 80.18 | 56,993 | 73.89 |
| ControlDay7Rep3 | 101,891 | 89,339 | 87.68 | 87,340 | 81,068 | 79.56 | 68,857 | 67.58 |
| ControlDay7Rep4 | 92,207 | 81,894 | 88.82 | 79,852 | 73,183 | 79.37 | 64,548 | 70 |
| ControlDay9Rep1 | 97,247 | 86,611 | 89.06 | 85,148 | 80,514 | 82.79 | 67,250 | 69.15 |
| ControlDay9Rep2 | 80,556 | 70,768 | 87.85 | 69,249 | 64,461 | 80.02 | 58,583 | 72.72 |
| ControlDay9Rep3 | 104,610 | 92,893 | 88.8 | 90,973 | 84,359 | 80.64 | 72,944 | 69.73 |
| ControlDay9Rep4 | 99,913 | 89,245 | 89.32 | 87,248 | 80,759 | 80.83 | 67,398 | 67.46 |

**Supplementary Information: Description 1:**

An overall significant difference in richness level (observed features) between all three groups was identified ($p = 0.011$). Pairwise comparisons identified richness level was significantly lower for the RInvT (241) group compared to the control (292) and pPNA (342) groups ($p = 0.011$, 0.016). However, richness level was not significantly different between control and pPNA groups ($p = 0.152$). The same trends between all three groups and pairwise comparisons did not change after rarefaction and / or filtering of chloroplast reads.

An overall significant difference between the relative evenness of the species' richness (Pielou's evenness) between the three groups ($p = 0.027$) was observed. Rarefaction did not greatly influence the outcome ($p = 0.031$). In contrast, filtering chloroplast did influence the outcome with a large increase in the level of non-significance ($p = 0.901$). Thus, removing chloroplast led to highly similar evenness between the species' richness of all three groups.

The diversity and richness accounting for both abundance and evenness of the taxa present (Shannon index) was significantly different between all three groups ($p = 0.018$). More specifically, the pPNA blocking protocol had a significantly higher average Shannon index (6.20) indicating more diverse species compared to the control (5.88) and RInvT (5.89) protocols ($p = 0.014$ and 0.015, respectively). This observation was not evident between control and the RInvT groups, indicating extremely similar species diversity levels ($p = 0.935$). The same results were achieved upon rarefaction. However, after filtering chloroplast reads, there was no significant difference between all three groups ($p = 0.05$). Control and pPNA method bacterial communities had similar species diversity (6.04 and 6.19, respectively) because the same hypervariable region was initially targeted, and chloroplast was then removed from control like pPNA blocking chloroplast leaving similar bacterial communities and species diversity ($p = 0.417$). Although group significance between all three groups was not significant, post-hoc pairwise comparisons continued to identify a significant difference between RInvT (5.88) and pPna groups ($p = 0.020$). In contrast to the unfiltered, rarefaction led to a different outcome where significant difference were identified between the Shannon index of all three groups ($p = 0.048$). However, pairwise comparisons only identified a significant difference between the RInvT and pPNA groups ($p = 0.016$).

Under all circumstances (i.e., filtering and rarefaction), the biodiversity measure that incorporates phylogenetic difference between species through the sum of length of branches (Faith's Phylogenetic Diversity) between all three groups was significantly different ($p < 0.05$). Both pPNA (21.39) and control (19.48) groups had significantly higher measures of biodiversity thus greater biodiversity compared to the RInvT (15.55) group ($p < 0.05$). Irrespective of whether filtering or rarefaction was applied, pPNA and control groups had similar biodiversity measures ($p = 0.358 – 0.417$).

**Supplementary Information: Table 2:** Alpha diversity metrics computed across time without chloroplast filtering using Qiime2 and without rarefaction. ± standard error. Letters a to d indicate significant differences across time between timepoints. *Indicates an overall significant effect across time. Letters A – B indicate significant differences between groups.

| **Observed features** | **Day 0** | **Day 2** | **Day 5** | **Day 7** | **Day 9** | |
|---|---|---|---|---|---|---|
| Control | 220.25 ± 49.85$^{a}$ | 344.75 ± 54.20$^{a}$ | 298.75 ± 6.142$^{a}$ | 339 ± 21.08$^{a}$ | 255 ± 20.29$^{a}$ | **AB** |
| BLOCK* | 398.25 ± 52.54$^{abc}$ | 525.75 ± 63.77$^{a}$ | 294.5 ± 52.43$^{bcd}$ | 329 ± 28.673$^{c}$ | 163 ± 15.40$^{d}$ | **A** |
| COMPETE* | 242 ± 33.81$^{a}$ | 312.50 ± 30.54$^{a}$ | 242.75 ± 13.19$^{a}$ | 225.75 ± 23.84$^{a}$ | 182 ± 12.77$^{a}$ | **B** |
| **Shannon** | **Day 0** | **Day 2** | **Day 5** | **Day 7** | **Day 9** | |
| Control* | 5.45 ± 0.16$^{a}$ | 6.19 ± 0.11$^{b}$ | 5.82 ± 0.09$^{ab}$ | 5.94 ± 0.18$^{ab}$ | 5.98 ± 0.15$^{ab}$ | **A** |
| BLOCK* | 6.13 ± 0.15$^{a}$ | 6.77 ± 0.16$^{b}$ | 6.18 ± 0.22$^{a}$ | 6.15 ± 0.14$^{a}$ | 5.79 ± 0.11$^{a}$ | **B** |
| COMPETE | 5.74 ± 0.13$^{a}$ | 6.18 ± 0.11$^{a}$ | 5.99 ± 0.05$^{a}$ | 5.78 ± 0.27$^{a}$ | 5.75 ± 0.12$^{a}$ | **A** |
| **FPD** | **Day 0** | **Day 2** | **Day 5** | **Day 7** | **Day 9** | |
| Control | 16.79 ± 3.28$^{a}$ | 22.88 ± 2.22$^{a}$ | 19.16 ± 1.29$^{a}$ | 22.68 ± 1.48$^{a}$ | 15.90 ± 1.49$^{a}$ | **A** |
| BLOCK* | 25.23 ± 2.33$^{abc}$ | 30.48 ± 2.80$^{a}$ | 18.50 ± 2.49$^{bcd}$ | 21.61 ± 1.79$^{c}$ | 11.12 ± 1.25$^{d}$ | **A** |
| COMPETE* | 17.22 ± 1.77$^{ab}$ | 20.23 ± 1.49$^{a}$ | 14.89 ± 1.28$^{ab}$ | 14.87 ± 0.32$^{ab}$ | 10.53 ± 0.96$^{b}$ | **B** |
| **Pielou's** | **Day 0** | **Day 2** | **Day 5** | **Day 7** | **Day 9** | |
| Control | 0.71 ± 0.02$^{a}$ | 0.74 ± 0.01$^{a}$ | 0.71 ± 0.01$^{a}$ | 0.71 ± 0.02$^{a}$ | 0.75 ± 0.01$^{a}$ | **A** |
| BLOCK* | 0.71 ± 0.02$^{a}$ | 0.75 ± 0.01$^{ab}$ | 0.76 ± 0.01$^{ab}$ | 0.74 ± 0.03$^{ab}$ | 0.79 ± 0.01$^{b}$ | **B** |
| COMPETE | 0.73 ± 0.01$^{a}$ | 0.75 ± 0.01$^{a}$ | 0.76 ± 0.01$^{a}$ | 0.74 ± 0.02$^{a}$ | 0.77 ± 0.01$^{a}$ | **B** |

**Supplementary Information: Table 3:** Alpha diversity metrics computed across time without chloroplast filtering using Qiime2 and with rarefaction. ± standard error. Letters a to d indicate significant differences across time between timepoints. *Indicates an overall significant effect across time. Letters A – B indicate significant differences between groups.

| **Observed features** | **Day 0** | **Day 2** | **Day 5** | **Day 7** | **Day 9** | |
|---|---|---|---|---|---|---|
| Control | 217.75 ± 48.53$^{a}$ | 335.5 ± 46.24$^{a}$ | 287.75 ± 6.47$^{a}$ | 326.5 ± 19.92$^{a}$ | 242.75 ± 18.56$^{a}$ | **AB** |
| BLOCK* | 381.5 ± 49.92$^{ab}$ | 507.25 ± 61.66$^{a}$ | 285.25 ± 48.98$^{ab}$ | 320.5 ± 26.80$^{ab}$ | 162 ± 15.56$^{b}$ | **A** |
| COMPETE* | 237.25 ± 32.42$^{a}$ | 302.25 ± 28.48$^{a}$ | 235.5 ± 10.44$^{a}$ | 219.25 ± 22.23$^{a}$ | 179.5 ± 12.29$^{a}$ | **B** |
| **Shannon** | **Day 0** | **Day 2** | **Day 5** | **Day 7** | **Day 9** | |
| Control* | 5.44 ± 0.16$^{a}$ | 6.20 ± 0.12$^{b}$ | 5.80 ± 0.09$^{ab}$ | 5.94 ± 0.17$^{ab}$ | 5.97 ± 0.15$^{ab}$ | **A** |
| BLOCK* | 6.13 ± 0.15$^{a}$ | 6.75 ± 0.16$^{b}$ | 6.16 ± 0.22$^{a}$ | 6.14 ± 0.14$^{a}$ | 5.79 ± 0.11$^{a}$ | **B** |
| COMPETE | 5.73 ± 0.13$^{a}$ | 6.17 ± 0.11$^{a}$ | 5.98 ± 0.05$^{a}$ | 5.77 ± 0.27$^{a}$ | 5.74 ± 0.12$^{a}$ | **A** |
| **FPD** | **Day 0** | **Day 2** | **Day 5** | **Day 7** | **Day 9** | |
| Control* | 16.53 ± 3.13$^{ab}$ | 22.13 ± 1.62$^{ab}$ | 17.80 ± 1.27$^{ab}$ | 21.44 ± 1.24$^{a}$ | 14.51 ± 1.32$^{b}$ | **A** |
| BLOCK* | 23.71 ± 2.15$^{ab}$ | 28.45 ± 2.51$^{a}$ | 17.76 ± 2.24$^{bc}$ | 20.68 ± 1.70$^{b}$ | 10.97 ± 1.31$^{c}$ | **A** |
| COMPETE* | 16.72 ± 1.63$^{ab}$ | 19.08 ± 1.35$^{a}$ | 13.90 ± 0.94$^{b}$ | 14.31 ± 0.25$^{bc}$ | 10.01 ± 0.87$^{d}$ | **B** |
| **Pielou's** | **Day 0** | **Day 2** | **Day 5** | **Day 7** | **Day 9** | |
| Control | 0.71 ± 0.02$^{a}$ | 0.74 ± 0.01$^{a}$ | 0.71 ± 0.01$^{a}$ | 0.71 ± 0.02$^{a}$ | 0.76 ± 0.01$^{a}$ | **A** |
| BLOCK | 0.72 ± 0.01$^{a}$ | 0.75 ± 0.01$^{ab}$ | 0.76 ± 0.01$^{ab}$ | 0.74 ± 0.03$^{ab}$ | 0.79 ± 0.01$^{b}$ | **B** |
| COMPETE | 0.73 ± 0.01$^{a}$ | 0.75 ± 0.01$^{a}$ | 0.76 ± 0.01$^{a}$ | 0.74 ± 0.02$^{a}$ | 0.77 ± 0.01$^{a}$ | **B** |

**Supplementary Information: Table 4:** Alpha diversity metrics computed across time with chloroplast filtering using Qiime2 but without rarefaction. ± standard error. Letters a to d indicate significant differences across time between timepoints. *Indicates an overall significant effect across time. Letters A – B indicate significant differences between groups.

| **Observed features** | **Day 0** | **Day 2** | **Day 5** | **Day 7** | **Day 9** | |
|---|---|---|---|---|---|---|
| Control | 216.25 ± 49.36$^{a}$ | 338.75 ± 52.86$^{a}$ | 294.25 ± 5.66$^{a}$ | 332.75 ± 21.00$^{a}$ | 251.50 ± 19.26$^{a}$ | **A** |
| BLOCK* | 391.75 ± 52.21$^{abc}$ | 514.75 ± 61.41$^{a}$ | 287.25 ± 50.97$^{bcd}$ | 322.35 ± 27.55$^{c}$ | 161.25 ± 14.87$^{d}$ | **A** |
| COMPETE* | 240 ± 33.58$^{a}$ | 310.25 ± 30.20$^{a}$ | 241 ± 13.46$^{a}$ | 224.25 ± 23.69$^{a}$ | 181.5 ± 12.65$^{a}$ | **B** |
| **Shannon** | **Day 0** | **Day 2** | **Day 5** | **Day 7** | **Day 9** | |
| Control* | 5.61 ± 0.17$^{a}$ | 6.35 ± 0.12$^{b}$ | 6.17 ± 0.06$^{ab}$ | 6.10 ± 0.19$^{ab}$ | 5.97 ± 0.15$^{ab}$ | **AB** |
| BLOCK* | 6.12 ± 0.15$^{a}$ | 6.75 ± 0.15$^{b}$ | 6.15 ± 0.21$^{a}$ | 6.14 ± 0.14$^{a}$ | 5.79 ± 0.16$^{a}$ | **A** |
| COMPETE | 5.73 ± 0.13$^{a}$ | 6.17 ± 0.11$^{a}$ | 5.99 ± 0.04$^{a}$ | 5.77 ± 0.27$^{a}$ | 5.74 ± 0.12$^{a}$ | **B** |
| **FPD** | **Day 0** | **Day 2** | **Day 5** | **Day 7** | **Day 9** | |
| Control | 17.08 ± 3.18$^{a}$ | 23.15 ± 2.10$^{a}$ | 19.98 ± 1.56$^{a}$ | 23.08 ± 1.70$^{a}$ | 16.13 ± 1.31$^{a}$ | **A** |
| BLOCK* | 25.33 ± 2.13$^{abc}$ | 30.69 ± 2.89$^{a}$ | 18.55 ± 21.71$^{bcd}$ | 21.71 ± 1.74$^{c}$ | 11.64 ± 1.22$^{d}$ | **A** |
| COMPETE* | 17.83 ± 1.77$^{ab}$ | 20.52 ± 1.46$^{a}$ | 15.01 ± 1.30$^{ab}$ | 15.65 ± 0.27$^{ab}$ | 11.04 ± 0.93$^{b}$ | **B** |
| **Pielou's** | **Day 0** | **Day 2** | **Day 5** | **Day 7** | **Day 9** | |
| Control | 0.74 ± 0.02$^{a}$ | 0.76 ± 0.01$^{a}$ | 0.75 ± 0.01$^{a}$ | 0.73 ± 0.02$^{a}$ | 0.75 ± 0.01$^{a}$ | **A** |
| BLOCK* | 0.71 ± 0.02$^{a}$ | 0.75 ± 0.01$^{ab}$ | 0.76 ± 0.01$^{ab}$ | 0.74 ± 0.03$^{ab}$ | 0.79 ± 0.01$^{b}$ | **A** |
| COMPETE | 0.73 ± 0.01$^{a}$ | 0.75 ± 0.01$^{a}$ | 0.76 ± 0.01$^{a}$ | 0.74 ± 0.03$^{a}$ | 0.77 ± 0.01$^{a}$ | **A** |

**Supplementary Information: Table 5:** Alpha diversity metrics computed across time with chloroplast filtering using Qiime2 and with rarefaction. ± standard error. Letters a to d indicate significant differences across time between timepoints. *Indicates an overall significant effect across time. Letters A – B indicate significant differences between groups.

| Observed features | Day 0 | Day 2 | Day 5 | Day 7 | Day 9 | |
|---|---|---|---|---|---|---|
| Control | 214.25 ± 48.38$^{a}$ | 329.75 ± 45.00$^{a}$ | 288.25 ± 6.61$^{a}$ | 317.5 ± 18.55$^{a}$ | 235.75 ± 17.53$^{a}$ | **AB** |
| BLOCK* | 373.5 ± 49.64$^{abc}$ | 490.75 ± 58.60$^{a}$ | 277.50 ± 47.63$^{bcd}$ | 312.50 ± 26.43$^{c}$ | 160.75 ± 14.61$^{d}$ | **A** |
| COMPETE | 231.25 ± 31.62$^{a}$ | 297 ± 28.89$^{a}$ | 232.5 ± 10.51$^{a}$ | 216 ± 21.39$^{a}$ | 179.25 ± 11.85$^{a}$ | **B** |
| **Shannon** | **Day 0** | **Day 2** | **Day 5** | **Day 7** | **Day 9** | |
| Control* | 5.62 ± 0.17$^{a}$ | 6.35 ± 0.12$^{b}$ | 6.16 ± 0.06$^{ab}$ | 6.09 ± 0.19$^{ab}$ | 5.95 ± 0.16$^{ab}$ | **AB** |
| BLOCK* | 6.10 ± 0.15$^{a}$ | 6.73 ± 0.15$^{b}$ | 6.14 ± 0.22$^{a}$ | 6.14 ± 0.14$^{a}$ | 5.79 ± 0.11$^{a}$ | **A** |
| COMPETE | 5.72 ± 0.13$^{a}$ | 6.16 ± 0.11$^{a}$ | 5.98 ± 0.05$^{a}$ | 5.76 ± 0.27$^{a}$ | 5.73 ± 0.12$^{a}$ | **B** |
| **FPD** | **Day 0** | **Day 2** | **Day 5** | **Day 7** | **Day 9** | |
| Control* | 16.90 ± 3.10$^{ab}$ | 22.39 ± 1.52$^{b}$ | 19.40 ± 1.65$^{ab}$ | 20.88 ± 0.65$^{b}$ | 14.66 ± 1.20$^{a}$ | **A** |
| BLOCK* | 23.70 ± 2.03$^{abc}$ | 28.09 ± 2.34$^{a}$ | 17.74 ± 2.20$^{bcd}$ | 20.61 ± 1.66$^{c}$ | 11.60 ± 1.21$^{d}$ | **A** |
| COMPETE* | 16.03 ± 1.93$^{ab}$ | 19.24 ± 1.42$^{b}$ | 13.95 ± 0.99$^{ab}$ | 14.77 ± 0.54$^{b}$ | 10.73 ± 0.85$^{a}$ | **B** |
| **Pielou's** | **Day 0** | **Day 2** | **Day 5** | **Day 7** | **Day 9** | |
| Control | 0.74 ± 0.02$^{a}$ | 0.76 ± 0.01$^{a}$ | 0.75 ± 0.01$^{a}$ | 0.73 ± 0.02$^{a}$ | 0.77 ± 0.01$^{a}$ | **A** |
| BLOCK* | 0.72 ± 0.01$^{a}$ | 0.76 ± 0.01$^{ab}$ | 0.77 ± 0.01$^{ab}$ | 0.74 ± 0.03$^{ab}$ | 0.79 ± 0.01$^{b}$ | **A** |
| COMPETE | 0.73 ± 0.01$^{a}$ | 0.75 ± 0.01$^{a}$ | 0.76 ± 0.01$^{a}$ | 0.75 ± 0.03$^{a}$ | 0.77 ± 0.01$^{a}$ | **A** |

**Supplementary Information: Table 6:** Significance of the overall effect of time on alpha diversity measures (RInvT, BLOCK and control measures grouped) as determined by linear mixed effects models.

| | Chloroplast not removed; Not rarefied | Chloroplast not removed; rarefied | Chloroplast removed; not rarefied | Chloroplast removed; rarefied |
|---|---|---|---|---|
| **Observed features** | 0.766 | 0.800 | 0.757 | 0.874 |
| **Faith's phylogenetic diversity** | 0.862 | 0.656 | 0.872 | 0.575 |
| **Shannon entropy** | 0.384 | 0.361 | 0.613 | 0.676 |
| **Pielou's evenness** | 0.580 | 0.401 | 0.867 | 0.976 |
